# Anatomically Programmable and Optically Tunable 3D-Bioprinted Tumor-Mimicking Phantoms for Quantitative Nanofluorophore Benchmarking

**DOI:** 10.64898/2026.10.18.757485

**Authors:** Itzel De Leon, Asma Harun, Indrajit Srivastava

## Abstract

Surgical resection remains a primary treatment for solid tumors, with fluorescence-guided surgery offering improved intraoperative visualization and precision. However, conventional murine models provide limited control over anatomical and optical variables critical to evaluating fluorescent imaging agents. Here, we developed 3D-bioprinted, anatomically tailored and optically tunable tumor-mimicking phantoms (TMPs) as a reproducible platform for quantitative nanofluorophore evaluation under tissue-relevant surgical conditions. These TMPs recreate subcutaneous, epithelial, and intraperitoneal tumor presentations while enabling controlled variation of lesion geometry, spatial distribution, tissue pigmentation, cellular composition, and nanofluorophore concentration. Semiconducting polymer nanofluorophores (SPNs) with distinct emission profiles were incorporated as model imaging agents and evaluated using an *in vivo* imaging system (IVIS). The engineered phantoms enabled quantitative assessment of wavelength-dependent fluorescence, lesion visualization and delineation across diverse geometries and tissue backgrounds, multiplexed imaging of spatially distinct lesions, and simulation of fluorescence-guided tumor resection. Constructs incorporating GFP-expressing cancer cells further demonstrated simultaneous visualization of cellular and nanoprobe-derived fluorescence. Collectively, these findings establish anatomically programmable TMPs as a versatile, reproducible framework for benchmarking fluorescent imaging agents, systematically interrogating anatomical and optical determinants of imaging performance, and advancing fluorescence-guided surgical strategies before *in vivo* evaluation and clinical translation.

## 1. Introduction

Accurate intraoperative identification of tumor margins remains a major challenge in the surgical management of tumors. Incomplete resection can leave residual malignant tissue and increase the likelihood of recurrence, whereas excessing removal of surrounding healthy tissue can compromise the function and quality of life. ^1, 2^ Fluorescence-guided surgery (FIGS) has emerged as a promising approach to address these challenges by providing real-time optical contrast between tumor and surrounding tissue during surgical procedures. ^3, 4^ Advances in fluorescent dyes, nanoparticles, and molecularly targeted imaging agents have expanded the repertoire of probes available for intraoperative tumor visualization.^4, 5^ The effectiveness of an optical probe depends not only on its intrinsic brightness but also on the anatomical and optical context in which it is used. ^6,7^ Factors such as tumor size, morphology, depth, spatial distribution, surrounding tissue composition, pigmentation, and wavelength-dependent absorption and scattering collectively influence fluorescence detection and tumor delineation. ^6, 8^

Systematic evaluation of these variables remains challenging with conventional preclinical models. ^9^ While animal models yield essential data on biodistribution, pharmacokinetics, targeting, toxicity, and biological response ^10^, individual anatomical and optical parameters are generally not amenable to independent manipulation. Variability in tumor implantation, growth, morphology, and burden among animals ^10, 11^ further complicates direct comparisons between imaging probes and hinders isolation of the effects of specific parameters on fluorescence performance.^8^ In contrast, tissue-mimicking phantoms offer defined, reproducible environments for investigating imaging systems and contrast mechanisms before applying them to more biologically complex models. ^6, 12^ Imaging phantoms have been employed for system calibration, comparison of imaging technologies, and quantitative evaluation under controlled experimental conditions.^8, 13^ Optical tissue phantoms also provide controlled settings to assess parameters such as imaging sensitivity, specificity, and spatial resolution, addressing the need for standardized platforms to benchmark emerging optical technologies.^6, 14, 15^

However, most existing platforms are designed primarily to replicate a specific tissue property, anatomical structure, or imaging modality, rather than to systematically evaluate fluorescent nanoprobes in tumor-specific environments. To address this gap, 3D tumor-mimicking phantoms (TMPs) were recently introduced by us as an intermediate platform for evaluating NIR-I/NIR-II/afterglow nanoprobes under conditions relevant to fluorescence-guided surgery. ^16–20^ These TMPs utilize gelatin-based matrices combined with hemoglobin and Intralipid to reproduce tissue absorption and scattering, while allowing precise control over nanoprobe concentration, and tumor-associated microenvironmental factors (pH, enzymes). This platform enables quantitative nanoprobe assessment, multiplexed fluorescence imaging, optical tissue-permeation profiling, tumor-margin delineation, and simulated image-guided surgery in *ex vivo* tissues. Notably, initial studies also demonstrated the feasibility of 3D printing TMPs with controlled and irregular geometries while maintaining reproducible material and fluorescence characteristics.

The initial framework primarily focused on replicating tumor optical properties and establishing TMPs as a platform for evaluating NIR nanoprobes under representative surgical conditions. A major remaining challenge is recreating the diversity of anatomical tumor presentations encountered in clinical practice while independently controlling lesion geometry, spatial organization, tissue background, and cellular composition.^16^ Tumors may develop as localized subcutaneous masses, irregular epithelial lesions, or multiple disseminated deposits within an anatomical cavity.^21,22^These variations can substantially influence the visualization and interpretation of fluorescent signals during surgery. A platform that can reproduce these distinct presentations would facilitate systematic investigation of how nanofluorophore properties interact with tumor anatomy and tissue optical context. This capability is particularly relevant for semiconducting polymer nanofluorophores (SPNs), whose optical properties can be engineered through polymer structure to yield nanoparticles with distinct excitation and emission profiles. Assessing how these spectral differences affect practical imaging performance requires evaluation beyond solution-based or homogeneous phantom matrices. Ideally, nanofluorophores should be assessed across controlled variations in lesion geometry, dimensions, spatial distribution, tissue optical background, nanoprobe concentration, and cellular composition, while maintaining reproducible experimental conditions. Additionally, such platforms should support procedures relevant to fluorescence-guided surgery, including tumor delineation, resection, residual-signal assessment, and multiplexed imaging.

This study presents anatomically tailored and optically tunable 3D-bioprinted TMPs for quantitative benchmarking of semiconducting polymer nanofluorophores. We incorporated three SPNs, with distinct fluorescence profiles into engineered tumor constructs representing subcutaneous, epithelial, and intraperitoneal disease scenarios (**Figure 1**). The platform allows precise modulation of lesion geometry, spatial distribution, tissue pigmentation, cellular composition, and nanofluorophore concentration while maintaining reproducible imaging conditions. These constructs enable investigation of wavelength-dependent nanoprobe visualization, tumor delineation within irregular tissue geometries, multiplexed detection of spectrally distinct lesions, and fluorescence retention during simulated tumor resection. Additionally, green fluorescent protein (GFP)-expressing ovarian cancer cells are incorporated to demonstrate independent visualization of cellular and nanoprobe-derived signals. Rather than replacing animal models, these 3D-bioprinted TMPs provide a controlled intermediate platform to prioritize nanofluorophores, dissect the contributions of anatomical and optical variables to imaging performance, and optimize fluorescence-guided surgical strategies before *in vivo* evaluation and clinical translation.

**Figure 1.**
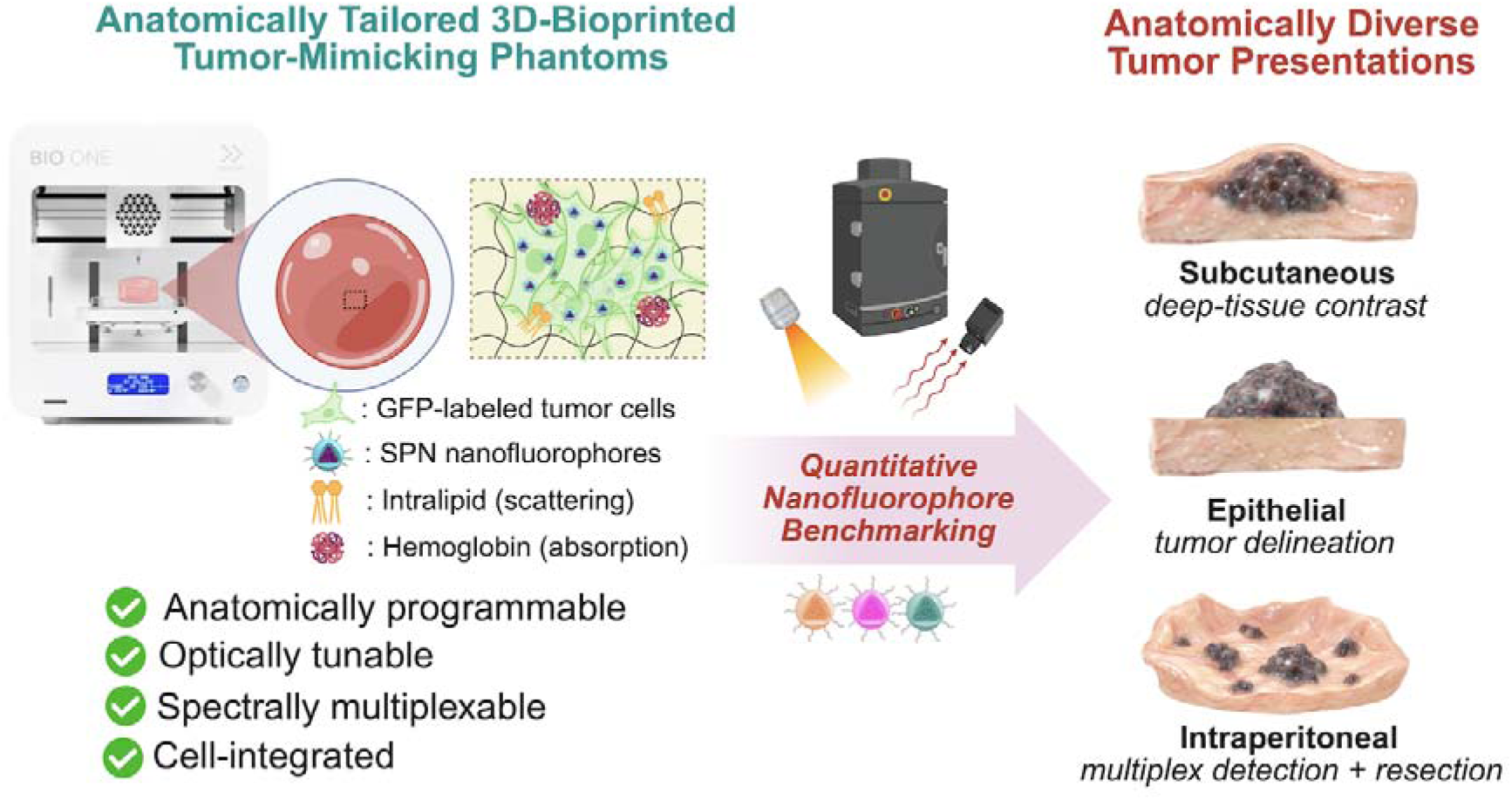
Schematic overview of anatomically tailored 3D-bioprinted TMPs for quantitative nanofluorophore benchmarking. The engineered TMP platform enables anatomical programmability, optical tunability, spectral multiplexing, and cellular integration. Three distinct tumor presentations, subcutaneous, epithelial, and intraperitoneal are modeled to evaluate tissue-dependent fluorescence contrast, tumor delineation, multiplexed imaging, and simulated fluorescence-guided surgical resection.

## 2. Results and Discussions

### 2.1. Synthesis and Characterization of the NIR-I Nanofluorophores

To evaluate how fluorescence wavelength influences nanoprobe performance within anatomically and optically complex tumor-mimicking phantoms, three semiconducting polymer with distinct optical properties were selected: poly[2-methoxy-5-(2-ethylhexyloxy)-1,4-phenylenevinylene] (MEHPPV), poly[2,7-(9,9-dioctylfluorene)-alt-4,7-bis(thiophen-2-yl)benzo-2,1,3-thiadiazole] (PFODBT), and poly[2,6-(4,4-bis-(2-ethylhexyl)-4H-cyclopenta [2,1-b;3,4-b′]dithiophene)-alt-4,7(2,1,3-benzothiadiazole)] (PCPDTBT). Each polymer was formulated into nanoparticles through nanoprecipitation using the amphiphilic phospholipid-polymer conjugate surfactant DSPE-PEG2K (**Figure 2a**). The resulting formulations are hereafter referred to MEHPPV NPs, PFODBT NPs and PCPDTBT NPs.

**Figure 2.**
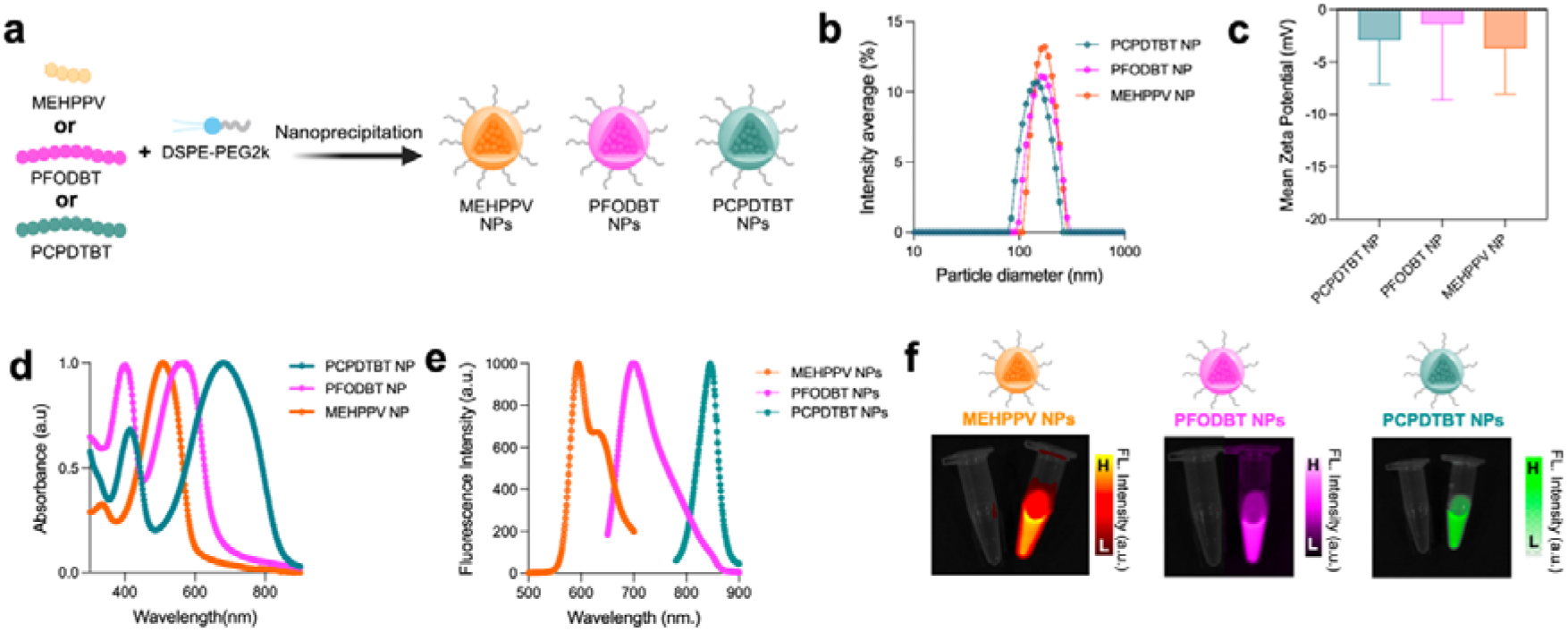
Preparation and physicochemical characterization of semiconducting polymer nanoparticles. (a) Schematic illustration of the preparation of MEHPPV, PFODBT and PCPDTBT NPs by DSPE-PEG2k-assisted nanoprecipitation. (b) Intensity-weighted hydrodynamic diameter distributions and (c) mean ζ-potential of three NPs measured by dynamic light scattering and electrophoretic light scattering, respectively. (d) Normalized UV-vis absorbance spectra and (e) fluorescence emission spectra demonstrating the distinct optical profiles of the NPs. (f) Representative IVIS fluorescence images acquired using excitation/emission filter pairs of 500 nm/ 580 nm for MEHPPV NPs, 605 nm/720 nm for PFODBT NPs, and 745 nm/840 nm for PCPDTBT NPs. Data in (c) represented as mean ± standard deviation, n = 3 independent prepared formulations.

Dynamic light scattering analysis revealed intensity-weighted hydrodynamic size diameters of approximately 195, 180, and 154 nm for MEHPPV, PFODBT, and PCPDTBT NPs, respectively (**Figure 2b**). The corresponding PDI values were 0.02, 0.2, and 0.10, indicating relatively narrow particle-size distribution and monodisperse populations. All three formulations exhibited moderately negative ζ-potentials (**Figure 2c**). Together with PEG-mediated steric stabilization, these surface characteristics supported the formation of well-dispersed nanoparticle suspensions (Table S1). Negative stained transmission electron microscope (TEM) imaging showed spherical morphology for every NP formulation **(Figure S3-S5)**. All three NP formulations maintained consistent hydrodynamic diameters and low PDI values over 7 days at 4°C, 37°C, and in plasma, demonstrating sustained colloidal stability without appreciable aggregation **(Figure S1-S2)**.

The nanoparticles retained the characteristic optical properties of their respective semiconducting polymer cores. MEHPPV, PFODBT, and PCPDTBT NPs exhibited an absorption maximum at 508, 572, and 680 nm, respectively (**Figure 2d**), with corresponding fluorescence emission maxima revealed emission maxima at 594, 697, and 846 nm (**Figure 2e**). ^16, 19, 23^. The dual absorption peaks observed in the UV-vis spectra of the three SPNs arise from distinct electronic transitions within their donor-acceptor conjugated polymer backbones. The higher energy peaks or shorter wavelength peaks are attributed to localized π-π* transitions of the donor segments, while the lower energy peaks or longer wavelength peaks originate from intramolecular charge transfer (ICT) interactions between donor and 7 acceptor moieties, resulting in extended absorption across the visible to NIR-I region.^24–27^ Consistent with the spectroscopic measurements, each formulation produced a strong fluorescence signal when imaged using its corresponding excitation and emission filters on the IVIS imaging system (**Figure 2f**). Collectively, these SPNs provide complementary emission profiles spanning the visible, far-red, and NIR-I regions, enabling systematic investigation of wavelength-dependent fluorescence behavior within the 3D-bioprinted tumor models.

### 2.2 Fluorescence Imaging Assessment of SPN Nanofluorophores in 3D-Printed Subcutaneous TMPs

Subcutaneous tumors develop beneath the skin and may arise from or extend into adipose connective and other soft tissues. In contrast to the irregular surface architecture of epithelial tumors, experimentally established subcutaneous lesions commonly exhibit smooth, rounded, or near-spherical geometries, although their dimensions and implantations depths can vary considerably.^11^ Lesion size, curvature, and tissue depth can substantially influence fluorescence attenuation and the apparent distribution of an imaging probe, making these features important when evaluating nanofluorophores for tumor detection and delineation. Although murine subcutaneous tumor models provide valuable biological information, they may require several weeks to establish and exhibit animal-to-animal differences in growth rate, size, shape, and implantation depth, limiting controlled comparisons of nanoprobe performance. ^28^

To reproduce these smooth, rounded morphology of subcutaneous lesions, the gelatin concentration and 3D printing parameters of the TMP bioink were optimized to provide sufficient flow during deposition while preserving structural integrity after printing. The reduced-viscosity formulation enabled controlled material deposition at higher printing velocities and volumetric flow rates. Printing into a rounded silicon mold followed by controlled cooling produced near-spherical tumor constructs. This approach enabled subcutaneous tumor models with predefined geometries and dimensions to be fabricated within approximately one hour, providing a rapid and reproducible platform for comparing the fluorescence performance of the three SPN formulations.

The fluorescence imaging performance of the three SPN formulations was evaluated using 3D-printed subcutaneous TMPs under tissue-mimicking conditions **(Figure 3a)**. The tumor constructs incorporated MEHPPV, PFODBT, or PCPDTBT NPs and were overlaid with porcine skin or muscle to simulate tissue-associated fluorescence attenuation **(Figure 3b)**. IVIS imaging was used to evaluate fluorescence distribution, tumor delineation, and tumor-to-normal tissue (T/NT) ratios for each formulation. All three SPN formulations produced detectable fluorescence signals through both porcine skin and muscle, with distinct differences in fluorescence intensity and tumor delineation. Under porcine skin, MEHPPV TMPs exhibited a relatively diffuse fluorescence distribution, with signal extending beyond the predefined tumor boundary. In contrast, PFODBT and PCPDTBT TMPs displayed more spatially confined fluorescence signals, allowing clearer visualization of the rounded tumor regions. Under porcine muscle, MEHPPV TMPs exhibited substantial fluorescence attenuation, resulting in limited contrast between the tumor and surrounding tissue. PFODBT and PCPDTBT TMPs maintained detectable fluorescence signals within the tumor regions, although the formulations showed differences in background fluorescence and signal distribution.

**Figure 3.**
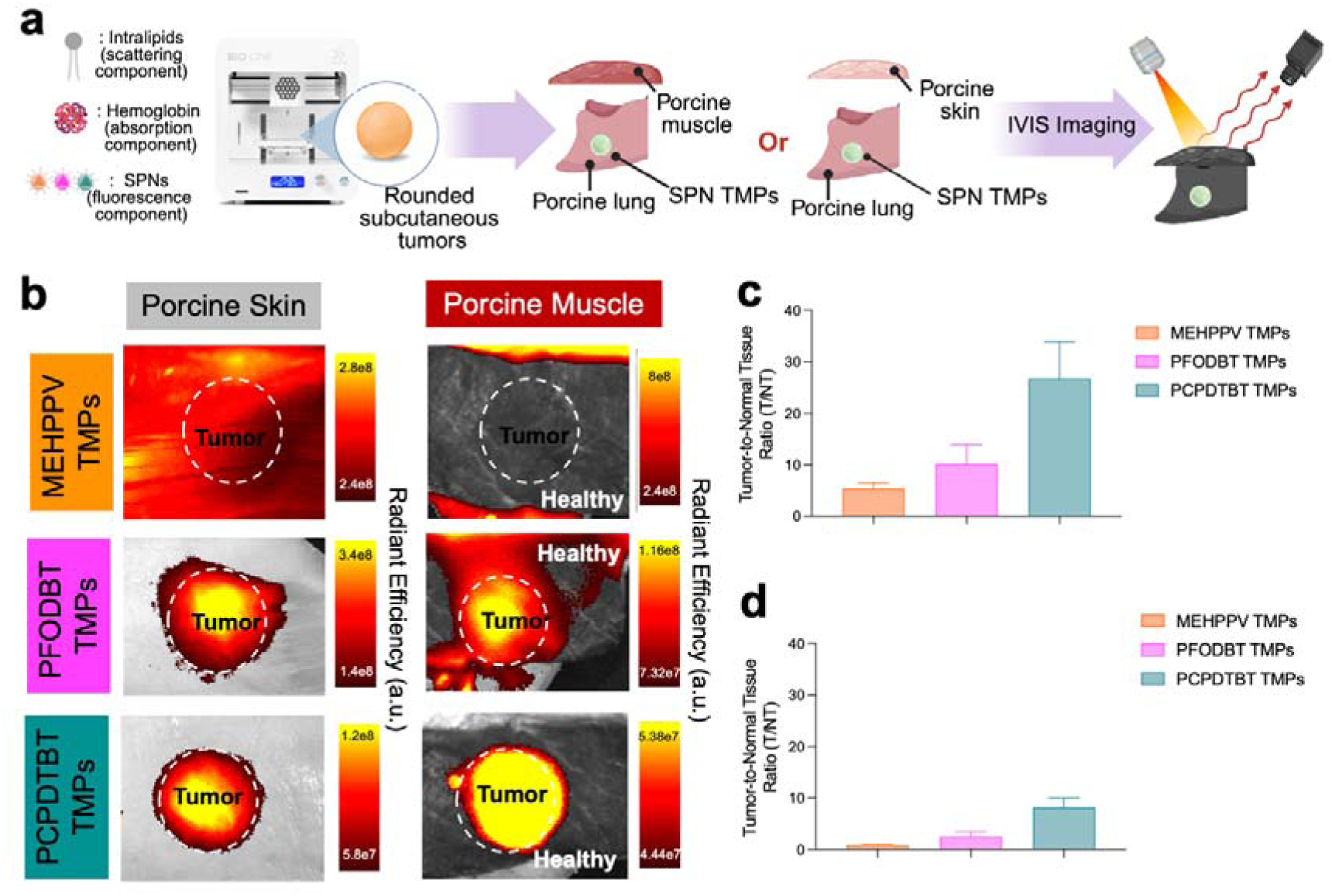
Fluorescence Imaging Assessment of SPN Nanofluorophores in 3D-Printed Subcutaneous TMPs. (a) Schematic illustrating the process of printing 3D-bioprinted subcutaneous tumor models and modeling using porcine lung, skin, and muscle within IVIS Imaging. (b) IVIS fluorescence imaging of the subcutaneous TMP with MEHPPV, PFODBT, and PCPDTBT under porcine skin and porcine muscle. (c-d) The area of healthy tissue as compared to tumor tissue was analyzed using IVIS imaging. As shown in IVIS, PCPDTBT seems to have the largest and most visible region between healthy and tumor tissue.

Quantitative analysis showed that the tumor to normal tissue ratios (T/NT ratios) were higher in porcine skin than in porcine muscle for all three SPN formulations **(Figure 3c, d)**. Among the tested formulations, PCPDTBT TMPs produced the highest fluorescence contrast in both tissue conditions, followed by PFODBT and MEHPPV TMPs. The lower T/NT ratios observed in porcine muscle may be attributed, at least in part, to differences in tissue optical properties, including absorption and scattering by endogenous tissue components. To further elucidate these effects, we evaluated the fluorescence responses of all three SPN formulations under varying concentrations of Intralipid (scattering component) and hemoglobin (absorption component). Increasing concentrations of these scattering and absorption agents attenuated fluorescence **(Figure S6-S11)**, supporting the role of tissue optical properties in modulating the detected fluorescence signal. Together, these findings highlight the influence of tissue composition on fluorescence contrast and establish the utility of 3D-printed subcutaneous TMPs as controlled platforms for evaluating nanofluorophore imaging performance across different tissue environments.

### 2.3 Fluorescence Imaging Assessment of SPN Nanofluorophores in 3D Printed Epithelial TMPs

Epithelial tumors arise from cells lining organ surfaces and cavities, with epithelial carcinomas accounting for the majority of ovarian malignancies.^29–31^ Unlike idealized spherical tumor models, epithelial ovarian tumors can exhibit irregular contours, curved surfaces, and spatially heterogeneous architectures.^32^ These morphological features can influence the apparent distribution and intensity of fluorescence by altering lesion thickness, imaging depth, surface orientation, and the optical path length between the imaging probes and detector. Reproducing this structural heterogeneity is therefore important when evaluating nanofluorophores intended for tumor delineation and fluorescence-guided surgical applications. Although murine epithelial tumor models^33, 34^ provide essential biological information, their irregular lesion morphology and optical heterogeneity are difficulty to control independently across animals, limiting systematic evaluation of how these features influence nanoprobe performance.

To reproduce the irregular contours and non-uniform surface architecture of epithelial tumors, the gelatin concentration and 3D-printing parameters of the TMP bioink were optimized to enable spatially variable material deposition while maintaining structural integrity following printing. The pre-flow volume, infill density, printing velocity, and volumetric extrusion rate were adjusted to generate constructs with curved surfaces and heterogeneous thicknesses (with the complete optimized printing parameters provided in the Methods section). Using the same parameters, a secondary TMP layer containing PCPDTBT nanofluorophores was deposited over the construct to create a fluorescent tumor-mimicking region across its irregular surface. The resulting epithelial TMPs were positioned on a synthetic skin mimic to reproduce a superficial tumor presentation. This approach produced epithelial TMPs with controlled but non-spherical morphologies, providing a reproducible platform for investigating how surface geometry, lesion thickness, and nanofluorophore distribution may influence fluorescence visualization and tumor delineation.

The fluorescence imaging performance of PCPDTBT NPs were subsequently evaluated using this engineered epithelial TMP platform to assess tumor visualization within irregular, tissue-analogous architectures **(Figure 4a)**. Two-dimensional fluorescence imaging demonstrated strong, localized emission from PCPDTBT-containing TMPs representing “tumor regions”, enabling visual differentiation from the surrounding tissue-mimicking background. Three-dimensional fluorescence reconstructions further demonstrated strong fluorescence localized within the epithelial tumor-mimicking regions, with comparatively lower fluorescence intensity in the underlying skin mimic, enabling clear visualization of the tumor-skin interface **(Figure 4b)**.

**Figure 4.**
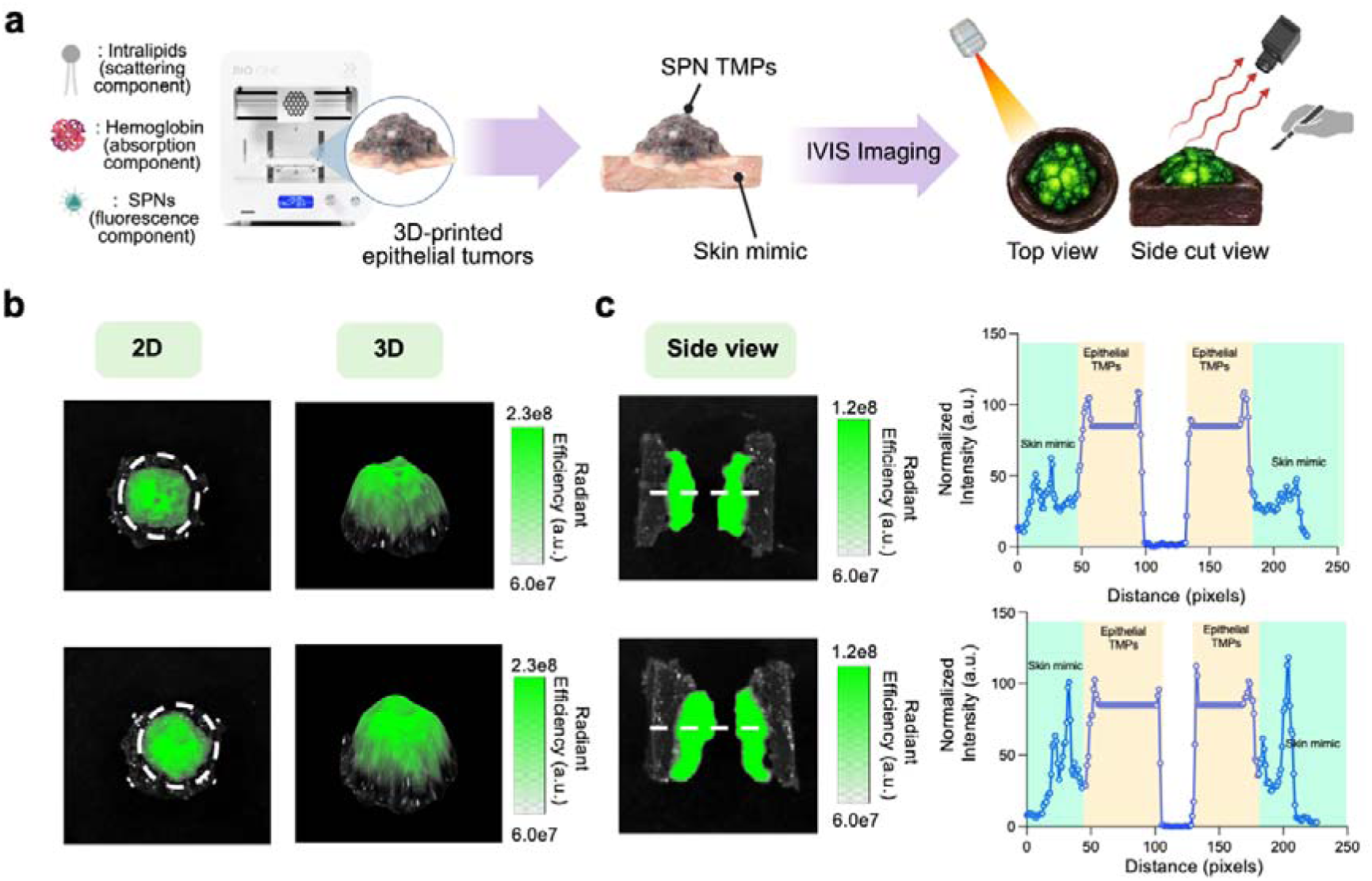
Optical characterization and spatial localization of SPN nanofluorophores in 3D-bioprinted epithelial TMPs. (a) Schematic illustrating the fabrication of epithelial TMPs incorporating SPNs as fluorescent components, hemoglobin as an absorption component, and Intralipid as a scattering component. The printed epithelial tumors are positioned on a synthetic skin mimic and evaluated using IVIS fluorescence imaging in top-view and side-cut configurations. (b) Representative 2D fluorescence images and corresponding 3D reconstructions demonstrating SPN fluorescence distribution within the epithelial TMPs using ImageJ. ^35^ (c) Representative side-view fluorescence images of sectioned epithelial TMPs and corresponding normalized fluorescence intensity line profiles acquired along the dashed lines, illustrating spatial localization of fluorescence within the tumor regions and their delineation from the surrounding skin mimic.

To evaluate fluorescence localization within the constructs, we sectioned the epithelial TMPs and underlying skin mimics to expose their cross-sectional architecture. Side-view imaging revealed localized fluorescence within the PCPDTBT-containing epithelial TMP regions, enabling visualization of their spatial distribution relative to the surrounding skin mimic (**Figure 4c**). The corresponding fluorescence intensity profiles exhibited elevated signals within the PCPDTBT-containing tumor regions, followed by pronounced reductions in intensity across the intervening nonfluorescent regions. Although measurable background fluorescence was observed within the surrounding skin mimic, the intensity differences enabled clear spatial differentiation between the fluorescent epithelial TMPs and adjacent tissue analogs. These findings demonstrate the utility of 3D-printed epithelial TMPs for evaluating nanoprobe fluorescence distribution and tumor delineation within geometrically complex tissue-mimicking environments.

### 2.4. Imaging-Guided Evaluation of SPN Nanofluorophores in 3D Bioprinted Intraperitoneal TMPs

Intraperitoneal tumor dissemination presents a challenging setting for fluorescence-guided imaging because lesions can occur at multiple anatomical locations and vary substantially in size, number, depth, and spatial distribution. Although murine intraperitoneal tumor models provide essential biological information and have been widely used to evaluate fluorescent nanoprobes ^36–38^, their implantation patterns and tumor burdens can vary among animals. Moreover, differences in anatomical dimensions and optical properties of nanofluorophores complicate controlled evaluation of how individual lesion characteristics influence nanofluorophores detection and surgical delineation. ^39–41^

Using this platform, we evaluated three complementary capabilities relevant to fluorescence-guided surgery: preservation of nanofluorophore fluorescence during simulated tumor resection, spectral differentiation of multiple SPN-containing lesions within a shared anatomical construct, and dual-channel visualization of SPNs and GFP-expressing tumor cells. Together, these studies demonstrate how anatomically tailored intraperitoneal phantoms can support controlled assessment of nanoprobe detection, multiplexed imaging, and simulated intraoperative performance.

#### 2.4.a. Simulated Fluorescence-Guided Resection of 3D Bioprinted Intraperitoneal TMPs

The 3D-printed intraperitoneal (IP) tumor model provides a spatially controlled platform to evaluate fluorescence-guided detection and resection of lesions containing MEHPPV, PFODBT, and PCPDTBT NPs. As illustrated in **Figure 5a**, tumor-mimicking lesions were precisely embedded within a melanin-containing tissue-mimetic matrix visually corresponding to approximately 4 on the skin color bar.^42^ Before resection, IVIS imaging revealed strong spatially localized fluorescence from each SPN-containing lesions with clear delineation from the surrounding background (**Figure 5b, d, f**).

**Figure 5.**
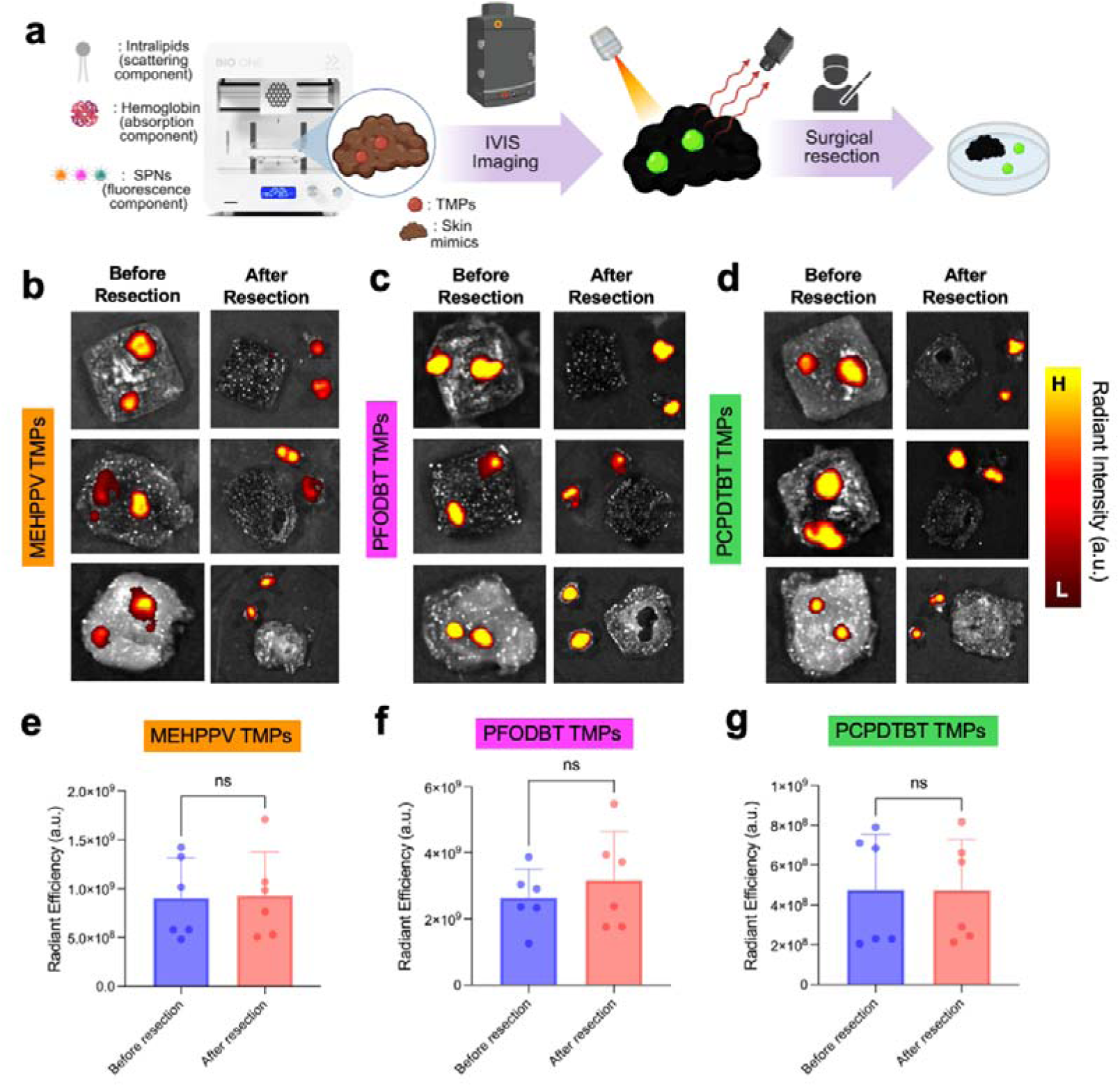
Simulated fluorescence-guided resection of SPN-containing intraperitoneal TMPs. (a) Schematic of intraperitoneal TMP fabrication, IVIS fluorescence imaging, and simulated surgical resection. Representative IVIS fluorescence images of TMPs consisting of (b) MEHPPV, (d) PFODBT and (f) PCPDTBT NPs positioned within a melanin-containing tissue-mimicking background before and after simulated surgical resection. Corresponding quantitative fluorescence analysis of lesions containing (c) MEHPPV, (e) PFODBT and (g) PCPDTBT NPs before and after resection. Data are presented as mean ± standard deviation (SD) (n=6 independently prepared phantoms); ns = not significant.

Following simulated surgical resection, the excised tumor constructs retained their fluorescence, with no statistically significant change in signal relative to their corresponding pre-resection measurements (**Figure 5c, e, g**). These findings indicate that the SPNs remained associated with the hydrogel constructs during surgical manipulation and that lesion fluorescence could be preserved after removal. The three formulations also enabled imaging through distinct spectral channels, demonstrating the adaptability of the intraperitoneal platform for evaluating nanofluorophores with different emission profiles. Collectively, these results establish the model as a reproducible system for assessing fluorescence-based lesion delineation and nanoprobe signal retention during simulated surgical resection.

#### 2.4.b. Multiplexed Nanoparticle Imaging in 3D-Printed Intraperitoneal Tumor Models

Fluorescence-guided surgery can improve tumor visualization and support the identification of residual disease during resection. However, most fluorescence-imaging approaches rely on a single contrast agent and therefore provide limited information about the molecular and spatial heterogeneity of tumors.^43, 44^ Multiplexed fluorescence imaging addresses this limitation by combining probes with distinguishable spectral signatures, enabling multiple tumor-associated features to be visualized within the same specimen.^45, 46^ When coupled with appropriate targeting ligands, such probes could be designed to recognize different cancer-cell populations, stromal components, immune-cell subsets, or molecular biomarkers within the tumor microenvironment. ^43, 47, 48^

Here, we evaluated whether MEHPPV, PFODBT, and PCPDTBT NPs, could be spatially distinguished within a single three-dimensional tissue-mimicking construct. This experiment was intended to establish a proof-of-concept platform for evaluating multiplexed fluorescence imaging under controlled, tissue-relevant optical conditions rather than to model biomarker-specific targeting. A melanin-containing tissue mimic was first printed using a concentrated porcine-gelatin formulation to reproduce tissue geometry and background autofluorescence.

Three spatially separated TMPs containing MEHPPV, PFODBT, or PCPDTBT NPs were subsequently printed onto the same tissue mimic by adjusting the printer’s *x*, *y*, and *z* coordinates (**Fig. 6a, S15**). The resulting construct was imaged using wavelength-specific excitation and emission settings selected for each SPN. Images of the melanin-containing background were also acquired to characterize and separate tissue-mimic autofluorescence. To investigate the influence of tissue-associated absorption on fluorescence performance, the effects of varying melanin concentrations on MEHPPV, PFODBT, and PCPDTBT NPs were evaluated. Increasing melanin concentrations resulted in fluorescence attenuation, highlighting the influence of melanin-mediated optical absorption on nanoprobe visualization **(Figure S12-S14)**.

**Figure 6.**
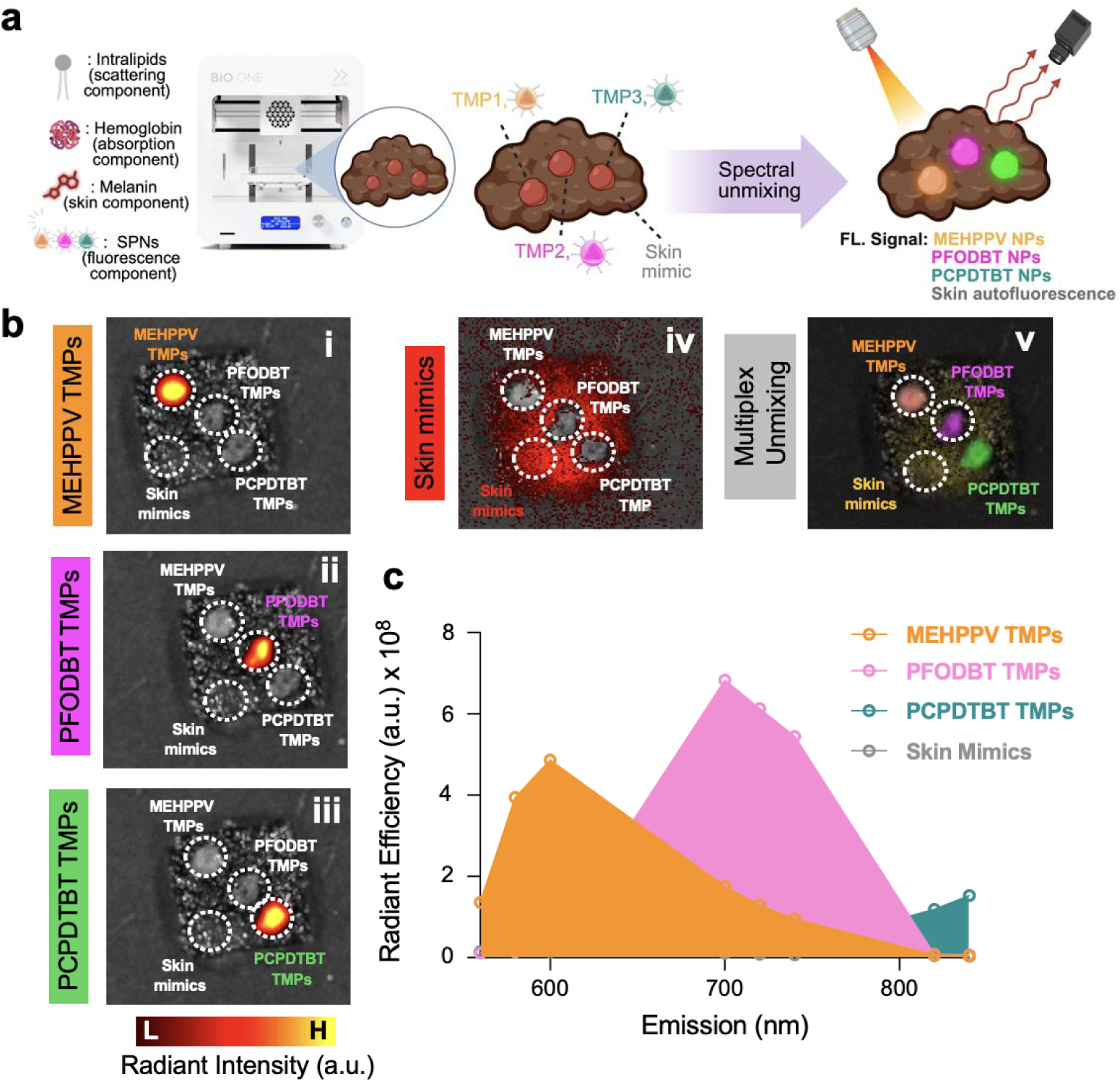
3D bioprinting of SPN-containing intraperitoneal TMPs for fluorescence multiplexing. (a) Schematic of the fabrication and spectral-unmixing workflow. Three spatially distinct TMPs containing MEHPPV, PFODBT, or PCPDTBT NPs were printed on to a single melanin-containing skin mimic. (b) Representative IVIS images acquired using wavelength-specific excitation and emission settings for (i) MEHPPV, (ii) PFODBT, and (iii) PCPDTBT TMPs; (iv) skin-mimic autofluorescence; and (v) the spectrally unmixed composite image. (c) Fluorescence emission profiles of the three SPN-containing TMPs and the skin mimic, illustrating their spectrally distinct signals used for multiplexed detection.

Each SPN-containing TMP produced a characteristic wavelength-dependent fluorescence response (**Fig. 6b, i-iii**). The melanin-containing tissue mimic exhibited a distinct background signal (**Fig. 6b, iv**), which was separated from the nanoparticle-associated signals during spectral unmixing. Combining these spectral profiles generated a composite image in which the three spatially distinct TMPs could be independently visualized within the same construct (**Fig. 6b, v**). The corresponding emission profiles further demonstrated differences among the MEHPPV-, PFODBT-, and PCPDTBT-containing TMPs and the tissue-mimicking background (**Fig. 6c**).

These findings demonstrate that the printed platform can incorporate multiple fluorescent nanoprobes and evaluate their spectral discrimination in a shared, optically complex environment. Although the present SPNs were used as spectrally distinct contrast agents rather than biomarker-specific probes, the platform could subsequently be adapted to test molecularly targeted nanoparticles designed to identify different tumor biomarkers or tissue compartments, as shown previously by us.^43^ It therefore provides a controlled approach for optimizing spectral separation, assessing interference from tissue autofluorescence, and developing multiplexed fluorescence-guided imaging strategies.

#### 2.4.c. Modeling Nanoprobe Accumulation in GFP-Expressing OVCAR3 Tumor Cells in Intraperitoneal TMPs

The preceeding studies evaluated the fluorescence behavior of SPNs within acellular tumor-mimicking constructs. To introduce a cellular component and distinguish tumor-associated cells from nanoprobe-derived fluorescence, GFP-expressing OVCAR3 ovarian cancer cells were incorporated into PCPDTBT NPs-containing intraperitoneal TMPs. GFP provided a secondary emission in the bioluminescence channel for confirming the presence and spatial location of tumor cells, while PCPDTBT NPs generated the primary NIR-I signal used for lesion visualization. This dual-channel design enabled independent optical detection of the cellular and nanoparticle components within the same tumor-mimicking constructs.

Three intraperitoneal TMPs with progressively decreasing lesion sizes were printed on a shared melanin-containing tissue-mimicking background (**Figure 7a**). The PCPDTBT NPs concentrations were decreased from 38 to 4 and 0.4 µg mL^−1^, while the corresponding GFP-expressing OVCAR3 cell densities were decreased from 6.5 × 10^5^ to 6.5 × 10^4^and 6.5 × 10^3^ cells mL^−1^, respectively **(Figure S16)**. By varying lesion size, nanoparticle concentration, and cellular density together, the constructs reproduced different levels of tumor and nanoprobe burden within a spatially controlled intraperitoneal model.

**Figure 7.**
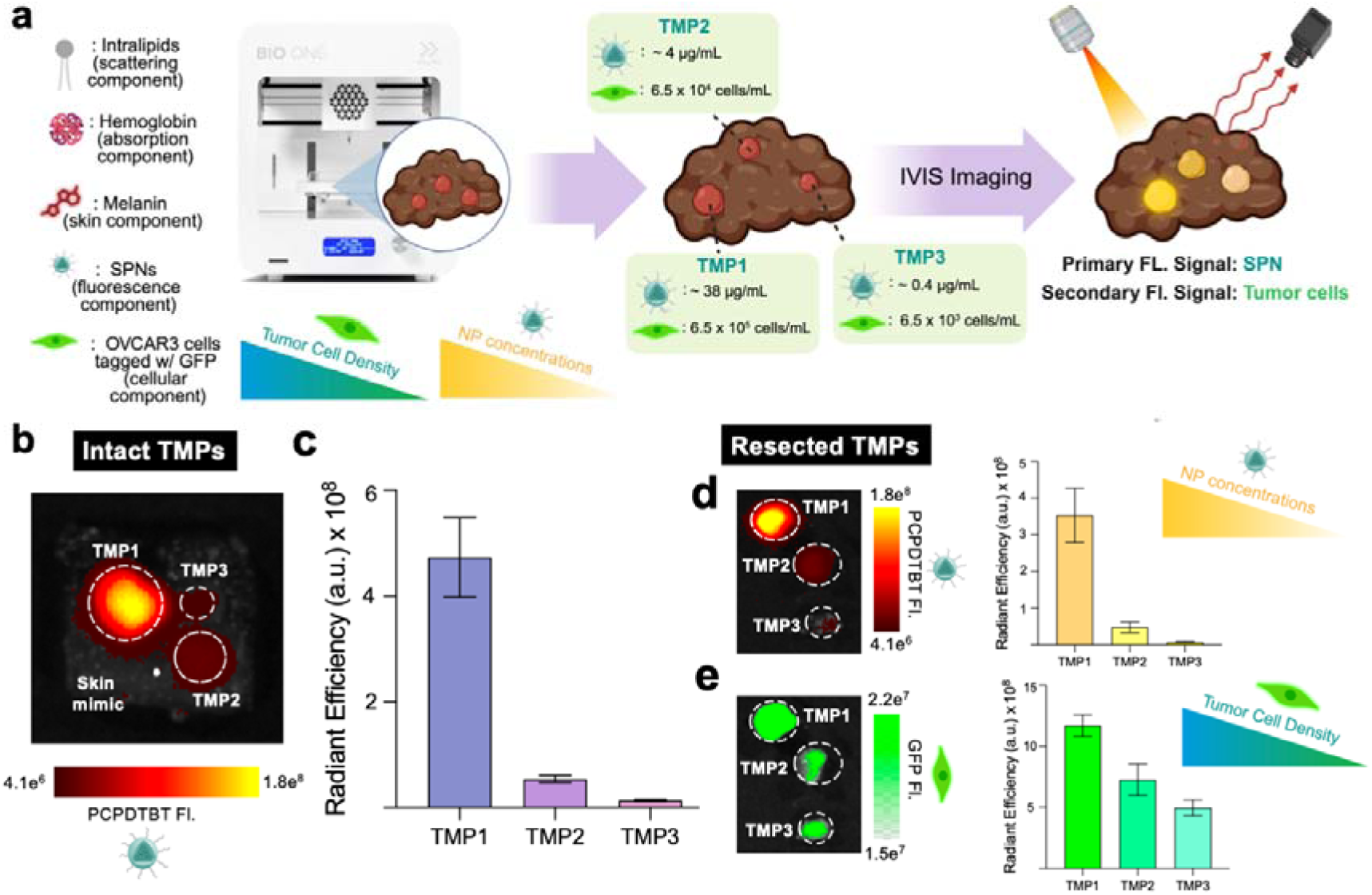
Dual-channel imaging of PCPDTBT NPs and GFP-expression OVCAR3 cells in 3D-Bioprinted Intraperitoneal TMPs. (a) Schematic illustrating the fabrication of intraperitoneal TMPs within varying lesion sizes, PCPDTBT NPs concentrations, and GFP-expressing OVCAR3 cell densities on a melanin-containing tissue-mimicking backgrounds. (b) Representative IVIS fluorescence image of the intact TMPs acquired in the quantitative analysis of radiant efficiency in the PCPDTBT channel using 745 nm excitation and 840 nm emission filters. (c) Corresponding quantitative analysis of radiant efficiency in the PCPDTBT channel. Representative fluorescence images of the resected TMPs and their corresponding radiant efficiency within (d) PCPDTBT NPs channel (Ex. 745 nm, Em. 840 nm) and (e) GFP channel (Ex. 465 nm, Em. 520 nm). Data are presented as mean ± standard deviation (n = 3 independently prepared phantoms).

IVIS imaging in the PCPDTBT channel clearly delineated the printed lesions from the surrounding tissue-mimicking background (**Figure 7b**). Quantitative analysis demonstrated progressively lower radiant efficiencies as lesion size and PCPDTBT NPs concentration decreased (**Figure 7c**). Because these variables were altered concurrently, the observed 20 differences represent their combined effects and cannot be attributed independently to lesion size or nanoparticle concentration. Following simulated resection, the excised lesions retained detectable PCPDTBT NPs fluorescence (**Figure 7d**), demonstrating that the nanoparticle-derived signal remained associated with the tumor-mimicking constructs during surgical manipulation. Imaging in the GFP channel independently confirmed the presence of OVCAR3 cells within the resected lesions (**Figure 7e**), although GFP fluorescence was weaker than the PCPDTBT NPs-derived signal under the respective imaging conditions. Collectively, these results demonstrate that the cellular and nanoprobe components of the intraperitoneal TMPs can be visualized through separate fluorescence channels, providing a controlled platform for evaluating tumor burden, nanoprobe-associated signal, and simulated fluorescence-guided resection.

## 3. Conclusion

We developed anatomically tailored, optically tunable 3D-bioprinted tumor-mimicking phantoms to evaluate semiconducting polymer nanofluorophores under controlled, tissue-relevant imaging conditions. We formulated MEHPPV, PFODBT, and PCPDTBT as colloidally stable nanoparticles with distinct emission profiles spanning the visible, far-red, and NIR-I regions. Incorporation of these nanofluorophores into epithelial, subcutaneous, and intraperitoneal TMPs enabled systematic assessment across tumor presentations with varying geometries, dimensions, spatial distributions, tissue backgrounds, and optical environments. In contrast to animal models, where these variables cannot be independently controlled, the printed constructs permitted reproducible definition of relevant anatomical and optical parameters, while supporting rapid fabrication and fluorescence analysis.

The intraperitoneal models demonstrated additional capabilities relevant to fluorescence-guided surgery. Lesions containing semiconducting polymer nanoparticles remained detectable after simulated resection, with minimal loss of fluorescence. Spectral unmixing enabled spatial discrimination of MEHPPV-, PFODBT-, and PCPDTBT-containing TMPs within a shared tissue-mimicking background. Incorporation of GFP-expressing OVCAR3 cells alongside PCPDTBT nanoparticles permitted independent visualization of cellular and nanoprobe-derived signals. These findings establish the platform as a versatile framework for benchmarking nanofluorophore performance, investigating the effects of tumor morphology and tissue optical properties, and optimizing multiplexed imaging and fluorescence-guided resection workflows. While these phantoms do not reproduce vascular perfusion, pharmacokinetics, or the full cellular complexity of living tumors, they complement conventional *in vitro* and animal models in a controlled way. Future integration of perfusable architectures, multiple cell populations, and molecularly targeted imaging probes could further expand their utility for prioritizing nanofluorophores and refining imaging strategies prior to *in vivo* evaluation and clinical translation.

## 4. Experimental Section/Methods

### Methods and Materials

DSPE-PEG2K was purchased from Biopharma PEG. All other chemicals were purchased from Sigma-Aldrich unless not stated otherwise.

### Preparation of MEHPPV NPs

To prepare MEHPPV NPs, 0.5 mg of MEHPPV polymer was co-dissolved with 20 mg of the surfactant DSPE-PEG2k in 1 mL of tetrahydrofuran (THF), serving as the organic phase. This solution was then added dropwise into 8 mL of 1x PBS aqueous phase in a 20 mL vial under probe sonication for 4 min. Subsequently, THF was removed by stirring the mixture at 950 rpm at room temperature overnight. The resulting nanoparticle suspension was purified by centrifugation using a 30 kDa molecular weight cut-off filter at 4500 rpm for 10 min, followed by three washes with 1x PBS to eliminate residual precursors. Finally, the suspension was passed through a 0.22-micron PVDF (polyvinylidene fluoride) filter, and the purified MEHPPV NPs were collected and stored at 4 °C for further use.

### Preparation of PFODBT NPs

To prepare PFODBT NPs, 0.5 mg of PFODBT polymer was co-dissolved with 5 mg of the surfactant DSPE-PEG2k in 2 mL of tetrahydrofuran (THF), serving as the organic phase. This solution was then added dropwise into 8 mL of 1x PBS aqueous phase in a 20 mL vial under probe sonication for 4 min. Subsequently, THF was removed by stirring the mixture at 950 rpm at room temperature overnight. The resulting nanoparticle suspension was purified by centrifugation using a 30 kDa molecular weight cut-off filter at 4500 rpm for 10 min, followed by three washes with 1x PBS to eliminate residual precursors. Finally, the suspension was passed through a 0.22-micron PVDF (polyvinylidene fluoride) filter, and the purified PFODBT NPs were collected and stored at 4 °C for further use.

### Preparation of PCPDTBT NPs

To prepare PCPDTBT NPs, 0.5 mg of PCPDTBT polymer was co-dissolved with 5 mg of the surfactant DSPE-PEG2k in 2 mL of tetrahydrofuran (THF), serving as the organic phase. This solution was then added dropwise into 8 mL of 1x PBS aqueous phase in a 20 mL vial under probe sonication for 4 min. Subsequently, THF was removed by stirring the mixture at 950 rpm at room temperature overnight. The resulting nanoparticle suspension was purified by centrifugation using a 30 kDa molecular weight cut-off filter at 4500 rpm for 10 min, followed by three washes with 1x PBS to eliminate residual precursors. Finally, the suspension was passed through a 0.22-micron PVDF (polyvinylidene fluoride) filter, and the purified PCPDTBT NPs were collected and stored at 4 °C for further use.

### Physicochemical and Optical Characterization of NPs

Dynamic light scattering (DLS) measurements were conducted to measure the hydrodynamic diameter of nanoparticles using a Litesizer DLS 700 Particle Analyzer instrument (Anton Paar). For each measurement, 50 μL of nanoparticle solution and 950 μL of PBS 1X were used. ζ-Potential measurements were also taken using the Litesizer DLS 700 Particle Analyzer instrument (Anton Paar). Ultraviolet–visible (UV–vis) absorbance was recorded on a Genesys 30 Visible Spectrophotometer (Thermo Fisher Scientific). The absorption spectra were collected at an interval of 1 nm scanning from 225 to 1100 nm. Fluorescence spectra were collected on an RF-6000 Spectro Fluorophotometer (Shimadzu). The emission spectra were collected at an interval of 1 nm scanning at an excitation wavelength of 490 nm and emission from 500 to 700 nm for MEHPPV NPs, at an excitation wavelength of 605 nm and emission from 650 to 900 nm for PFODBT NPs, and at an excitation wavelength of 645 nm and emission from 780 to 900 nm for PCPDTBT NPs. Fluorescence imaging was conducted in the IVIS imager, at an excitation filter of 500 nm and an emission filter of 580 nm for MEHPPV NPs, an excitation filter of 605 nm and emission filter of 720 nm for PFODBT NPs, and an excitation filter of 745 nm and emission filter of 840 nm for PCPDTBT NPs.

### 3D Bioprinting of Tumor-Mimicking Phantoms

Anatomically distinct TMPs, representing epithelial, subcutaneous, and intraperitoneal tumor presentations, were fabricated using a BIO ONE 3D bioprinter (CELLINK). Gelatin-based bioinks incorporating hemoglobin and Intralipid as optical absorption and scattering components, respectively, were formulated and optimized for each anatomical model. Bioink composition and printing parameters were adjusted to reproduce the characteristic geometries and spatial distributions of the different tumor presentations, as detailed in the following subsections.

### 3D Bio-Printing of Epithelial Tumors

Epithelial tumor-mimicking phantoms (TMPs) were fabricated using a high-concentration gelatin-based bioink to reproduce the irregular surface morphology and three-dimensional architecture of epithelial ovarian tumors. The bioink was prepared by combining Intralipid (900 µL), hemoglobin (150 mg), and gelatin (1020 mg) in 17 mL of PBS 1x. Hemoglobin and Intralipid served as the optical absorption and scattering components, respectively, to mimic the optical properties of biological tissues. The prepared bioink was transferred into a syringe compatible with the 3D bioprinter and deposited using optimized printing parameters, including a pre-flow volume of 25 µL, extrusion volume of 10 µL, and printing speed of 10 mm s^−1^. These parameters facilitated the fabrication of epithelial TMPs with irregular surface contours while maintaining their structural integrity.

To generate nanofluorophore-containing epithelial tumor regions, a secondary bioink was prepared by mixing 1.5 mL of the gelatin-based TMP formulation with 0.5 mL of PCPDTBT nanoparticle suspension. The resulting formulation was transferred into a separate printing syringe and deposited as a fluorescent top layer over the previously printed epithelial TMPs using the same printing parameters. This sequential bioprinting approach enabled the incorporation of PCPDTBT nanofluorophores within anatomically irregular tumor-mimicking constructs for subsequent fluorescence imaging and tumor delineation studies.

### 3D Bio-Printing of Subcutaneous Tumors

Subcutaneous TMPs were fabricated using a gelatin-based bioink optimized to produce smooth, rounded tumor geometries representative of subcutaneous lesions. The bioink contained gelatin (720 mg) and the previously described hemoglobin-and Intralipid-containing tissue-mimicking components in PBS 1x. Compared with the higher gelatin concentration used for epithelial constructs, the reduced gelatin content was selected to facilitate extrusion and the formation of rounded geometries. For nanofluorophore incorporation, 2 mL of the prepared TMP bioink was mixed with 0.5 mL of a 100-µg mL^−1^ suspension of MEHPPV, PFODBT, or PCPDTBT nanoparticles. Each nanofluorophore formulation was prepared separately to generate distinct fluorescent tumor constructs.

The nanoparticle-containing bioinks were transferred into printing syringes and briefly cooled at −20°C before bioprinting. Printing parameters were adjusted to promote uniform deposition and preserve the rounded morphology of the resulting constructs. The TMPs were formed using rounded molds to obtain reproducible, near-spherical geometries. To establish a tissue-associated imaging configuration, appropriately sized cavities were prepared in excised porcine lung tissue, into which the fabricated TMPs were placed. The constructs were subsequently evaluated under porcine skin and muscle tissue to investigate tissue-dependent fluorescence attenuation, tumor delineation, and nanofluorophore imaging performance.

### 3D Bio-Printing of Intraperitoneal Tumor Phantoms

Intraperitoneal TMPs were fabricated using a sequential 3D bioprinting approach to reproduce spatially distributed tumor lesions within a tissue-mimicking background. A melanin-containing bioink was first prepared by combining 500 µL of melanin solution (2.5 µg mL^−1^) with 1.5 mL of a 6% gelatin formulation. Melanin was incorporated to introduce tissue-associated pigmentation and optical absorption. The resulting formulation was transferred into a printing syringe and deposited to generate a flat, skin-like tissue-mimicking substrate.

Tumor-mimicking lesions were subsequently printed onto the prepared substrate using a gelatin-based bioink containing Intralipid (900 µL), hemoglobin (150 mg), and gelatin (720 mg) in 17 mL of PBS 1x. The bioink was loaded into the printer syringe and deposited at predefined locations by controlling the x-, y-, and z-axis translations of the printing nozzle. Individual tumor features were fabricated using a nominal printing scale of 1 mm x 1 mm x 0.1 mm. The printing parameters were adapted from those used for the tissue-mimicking substrate, with an increased pre-flow volume to facilitate deposition of the tumor-containing bioink. This sequential fabrication approach enabled controlled spatial positioning of multiple tumor-mimicking lesions within a shared tissue-mimicking construct. Nanofluorophore-containing TMPs incorporating MEHPPV, PFODBT, or PCPDTBT nanoparticles were subsequently used for fluorescence imaging, spectral discrimination, and simulated fluorescence-guided surgical resection.

### OVCAR3 Cells Culture

GFP-tagged OVCAR3 cells, a human ovarian adenocarcinoma cell line, were cultured in RPMI 1640 medium supplemented with 10% FBS, 1% penicillin-streptomycin, and 1% Antibiotic-Antimycotic. When the cells reached 70% confluency, they were detached, centrifuged, and resuspended in PBS1X.

## Acknowledgements.

This work was supported by grants from the Cancer Prevention and Research Institute of Texas (RP260870 to I.S.), the National Institutes of Health (R35GM166367 to I.S.), and American Heart Association (26CDA1597763 to I.S.). The authors acknowledge Mahenour Megahed for OVCAR3 cell culture, Elizabeth Chong for colloidal stability measurement, and Md. Hasnat Rashid for TEM imaging. All schematic figures were made in Biorender by I.S., except parts of Figure 4a scheme were generated with ChatGPT.

## Author Contributions

I. S. conceived the idea and design the experiments. I.D.L. and A.H. led the overall study. 3D bioprinting of TMPs were conducted by I. D. L. under the supervision of I.S. A.H. synthesized and characterized the SPN nanofluorophores under the supervision of I.S. A.H. and I.D.L. performed IVIS data collection and analysis under the supervision of I.S. The first draft of the manuscript was written by I.D.L. and A.H., with edits from I.S. All authors have approved the final version of the manuscript.

