## Supporting Information for "Anatomically Programmable and Optically Tunable 3D-Bioprinted Tumor-Mimicking Phantoms for Quantitative Nanofluorophore Benchmarking"

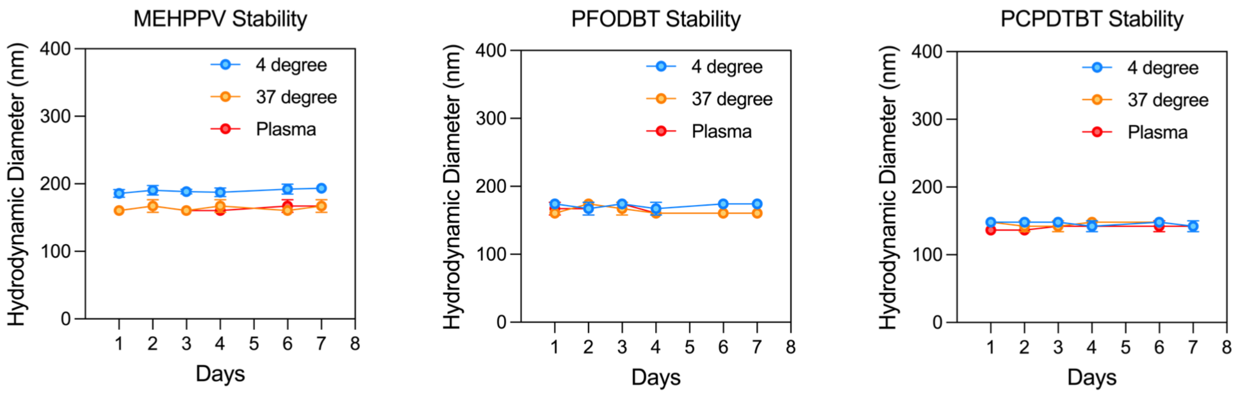


**Figure S1.** Hydrodynamic size stability of the NPs under different storage conditions over a 7 days period.

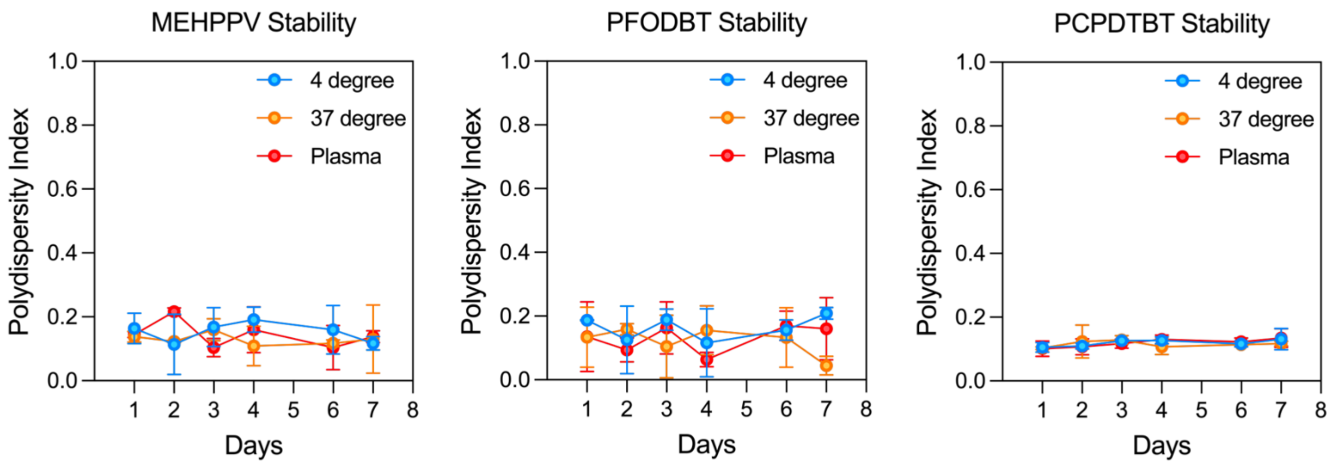


**Figure S2.** Polydispersity indices of the NPs under different storage conditions over a period of 7 days.

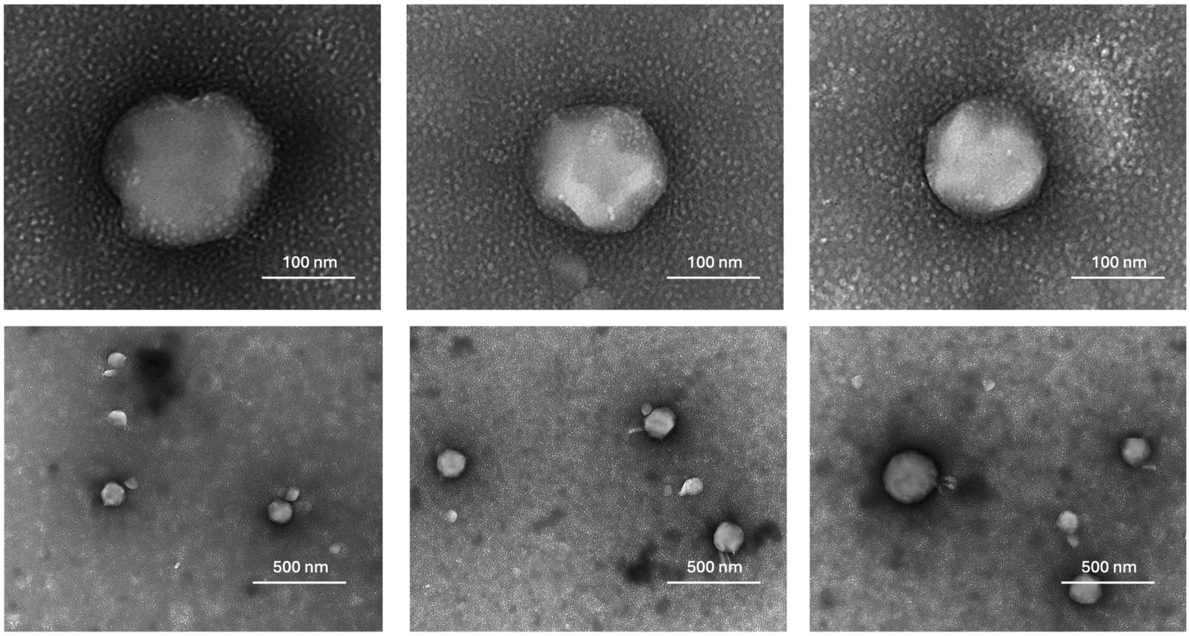


**Figure S3.** Negative-stained TEM images of MEHPPV NPs. Scale bars are indicated in each image.

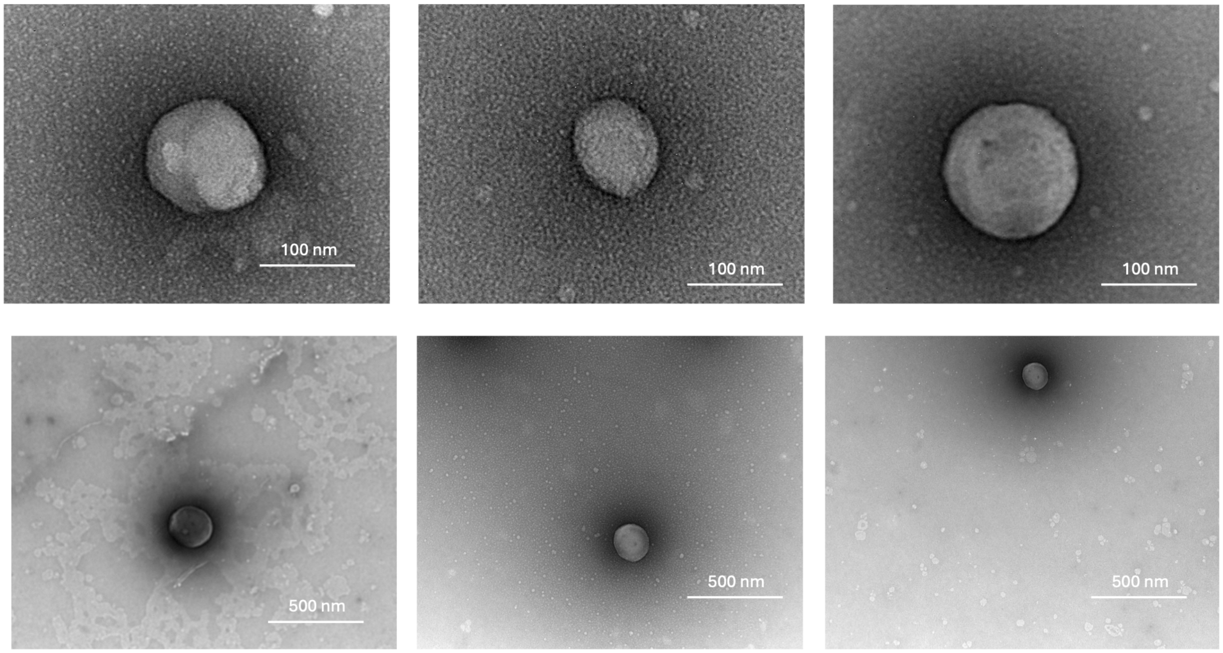


**Figure S4.** Negative-stained TEM images of PFODBT NPs. Scale bars are indicated in each image.

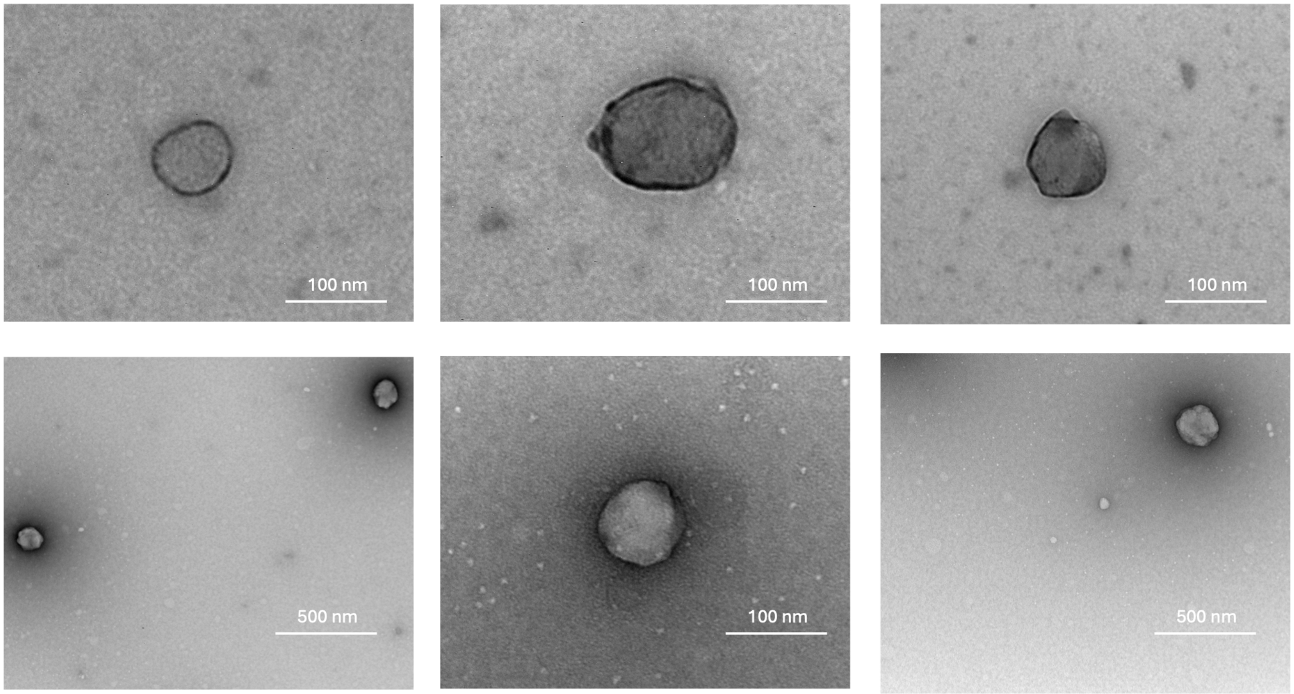


**Figure S5.** Negative-stained TEM images of PCPDTBT NPs. Scale bars are indicated in each image.

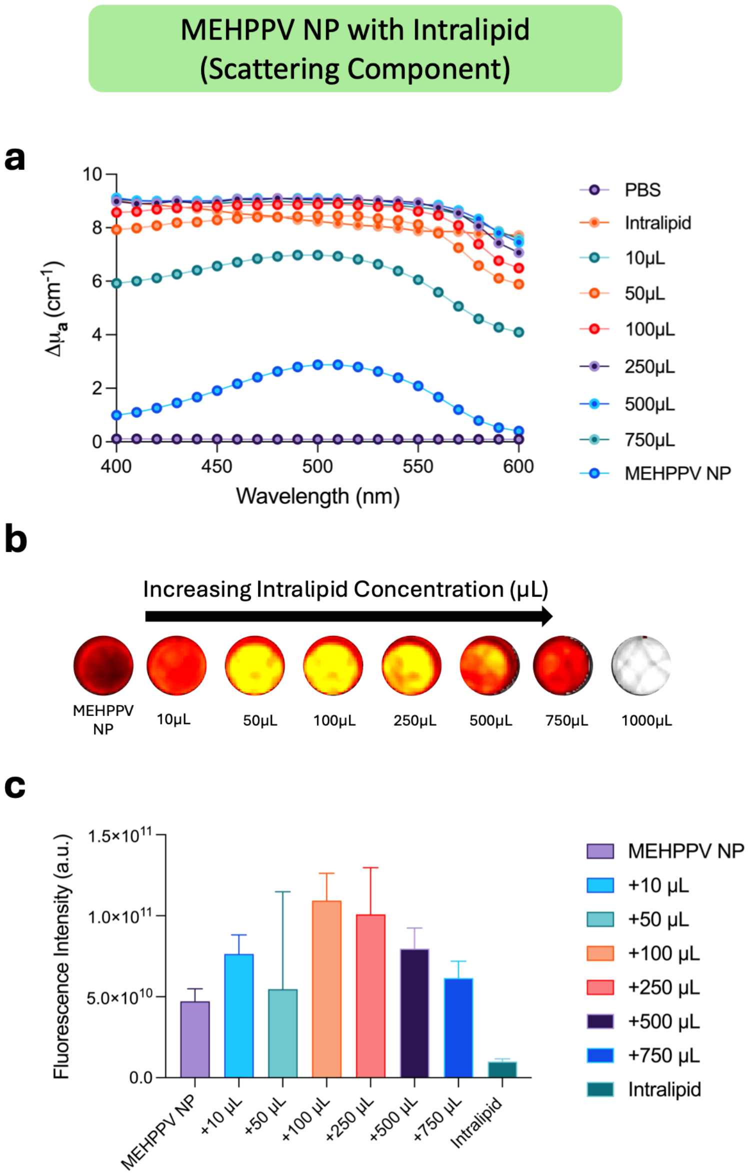


**Figure S6. Effect of Intralipid concentration on the fluorescence of MEHPPV NPs.** (a) Changes in the absorption coefficient ($\Delta\mu_a$) of MEHPPV NPs with increasing volumes of Intralipid. (b) Representative fluorescence images of MEHPPV NPs in the presence of increasing volumes of Intralipid (10–1000 μL). (c) Quantitative analysis of fluorescence intensity as a function of added Intralipid volume. Data are presented as mean ± SD.

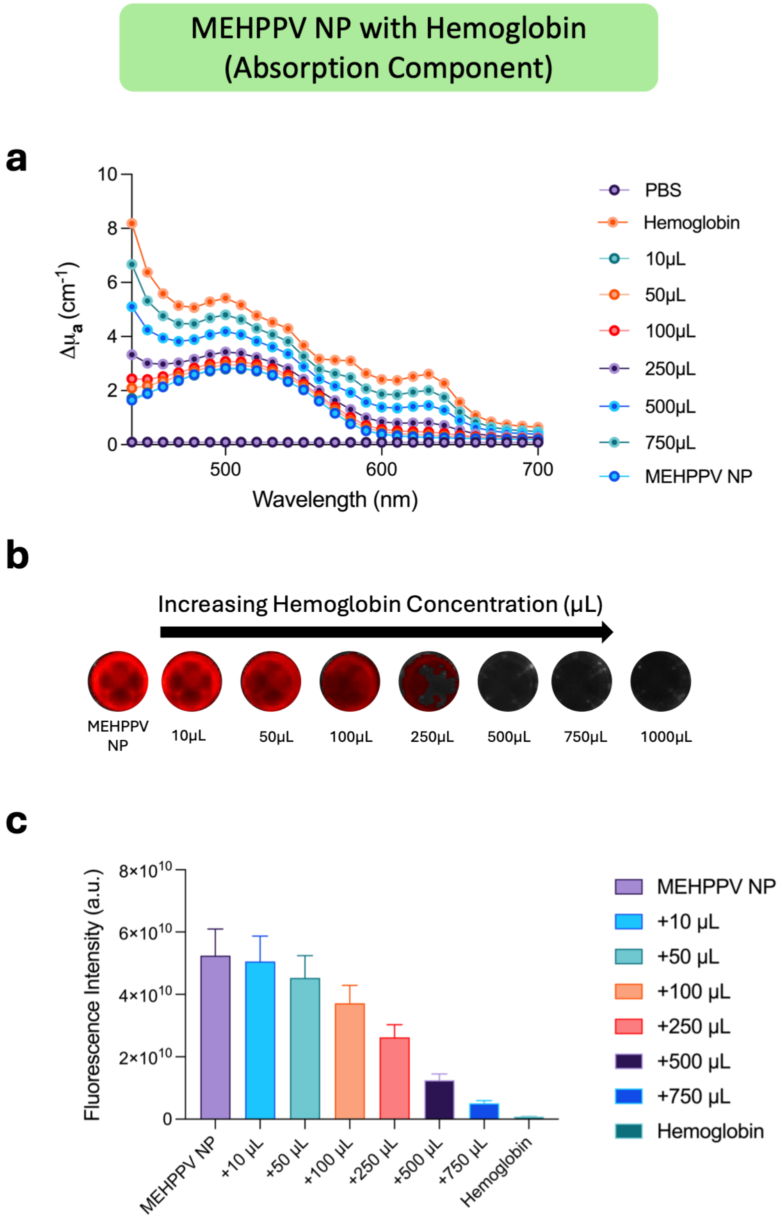


**Figure S7. Effect of Hemoglobin concentration on the fluorescence of MEHPPV NPs.** (a) Changes in the absorption coefficient ($\Delta\mu_a$) of MEHPPV NPs with increasing volumes of hemoglobin. (b) Representative fluorescence images of MEHPPV NPs in the presence of increasing volumes of hemoglobin volumes (10–1000 μL). (c) Quantitative analysis of fluorescence intensity as a function of added hemoglobin volume. Data are presented as mean ± SD.

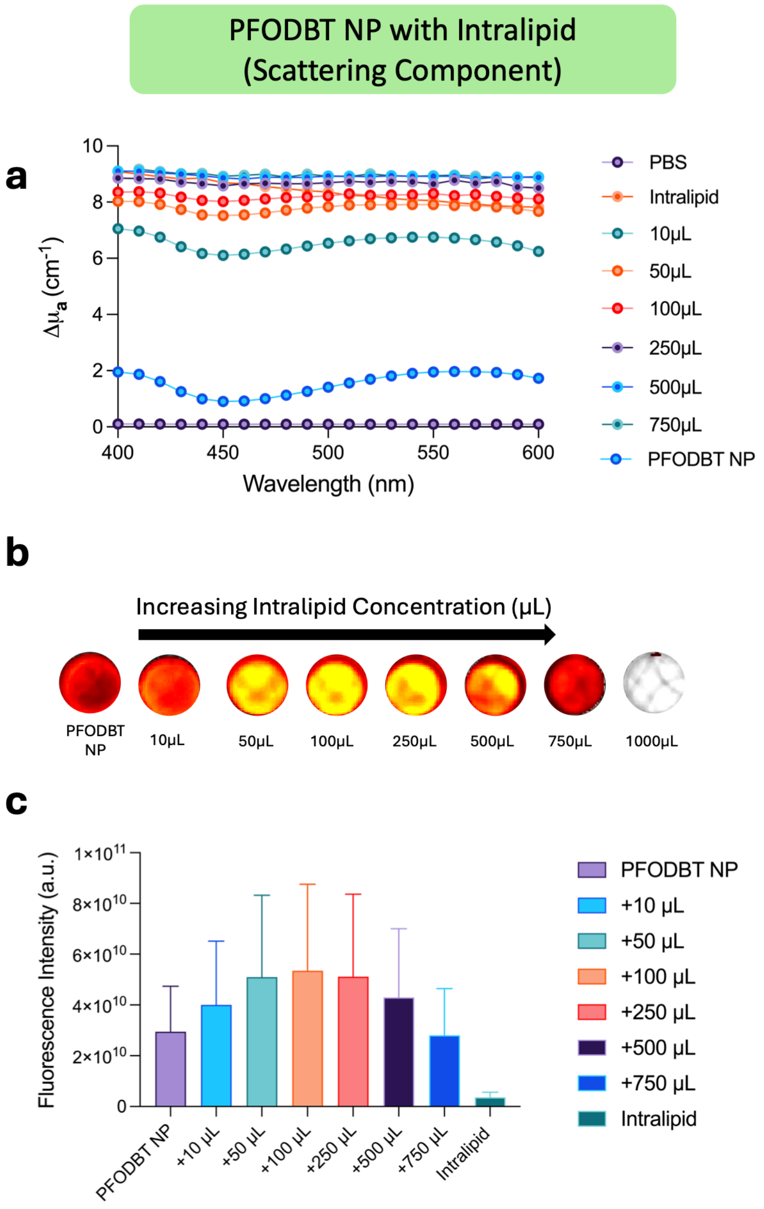


**Figure S8. Effect of the added volume of Intralipid on the fluorescence of PFODBT NPs.** (a) Changes in the absorption coefficient ($\Delta\mu_a$) of PFODBT NPs with increasing volumes of Intralipid. (b) Representative fluorescence images of PFODBT NPs in the presence of increasing volumes of Intralipid (10-1000 μL). (c) Quantitative analysis of fluorescence intensity as a function of added Intralipid volume. Data are presented as mean ± SD.

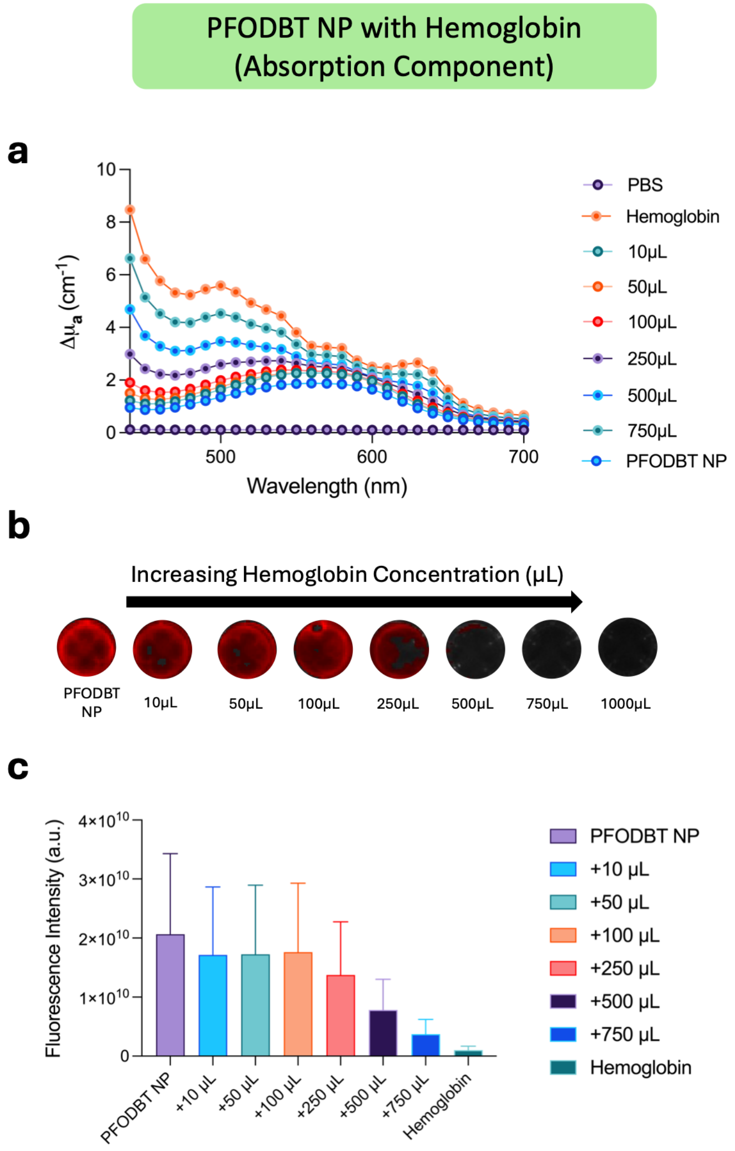


**Figure S9. Effect of the added volume of hemoglobin on the fluorescence of PFODBT NPs.** (a) Changes in the absorption coefficient ($\Delta\mu_a$) of PFODBT NPs with increasing volumes of hemoglobin. (b) Representative fluorescence images of PFODBT NPs in the presence of increasing volumes of hemoglobin (10-1000 μL). (c) Quantitative analysis of fluorescence intensity as a function of added hemoglobin volume. Data are presented as mean ± SD.

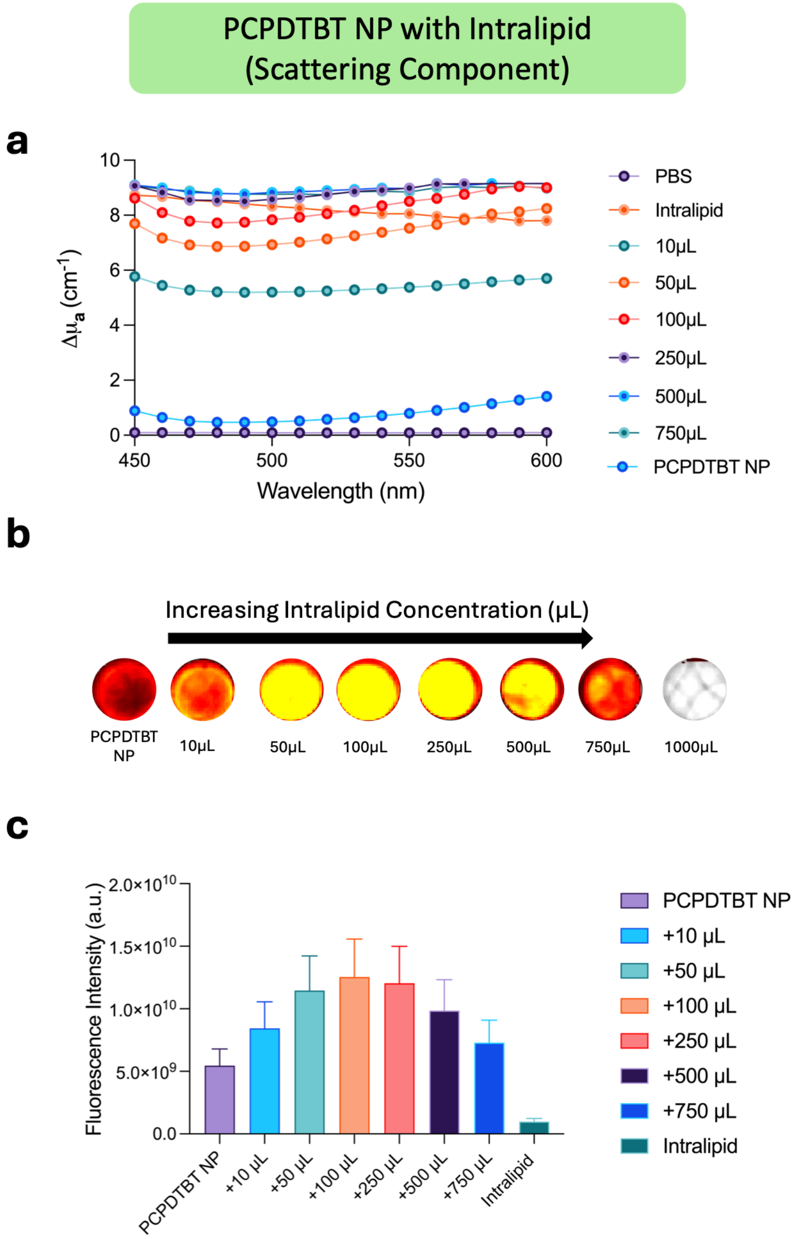


**Figure S10. Effect of the added volume of Intralipid on the fluorescence of PCPDTBT NPs.** (a) Changes in the absorption coefficient ($\Delta\mu_a$) of PCPDTBT NPs with increasing volumes of Intralipid. (b) Representative fluorescence images of PCPDTBT NPs in the presence of increasing volumes of Intralipid (10-1000 μL). (c) Quantitative analysis of fluorescence intensity as a function of added Intralipid volume. Data are presented as mean ± SD.

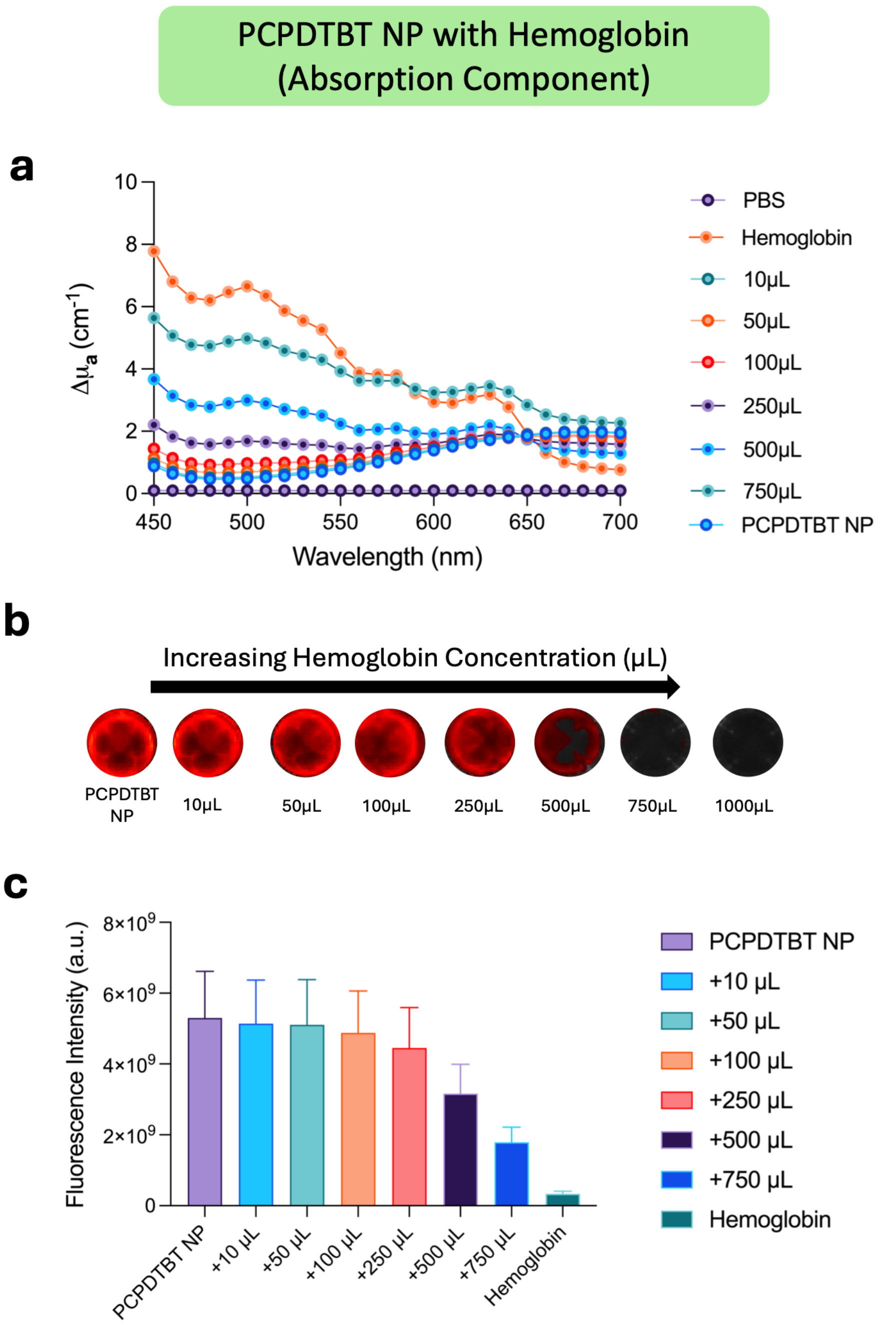


**Figure S11. Effect of the added volume of hemoglobin on the fluorescence of PCPDTBT NPs.** (a) Changes in the absorption coefficient ($\Delta\mu_a$) of PCPDTBT NPs with increasing volumes of hemoglobin. (b) Representative fluorescence images of PCPDTBT NPs in the presence of increasing volumes of hemoglobin (10-1000 μL). (c) Quantitative analysis of fluorescence intensity as a function of added hemoglobin volume. Data are presented as mean ± SD.

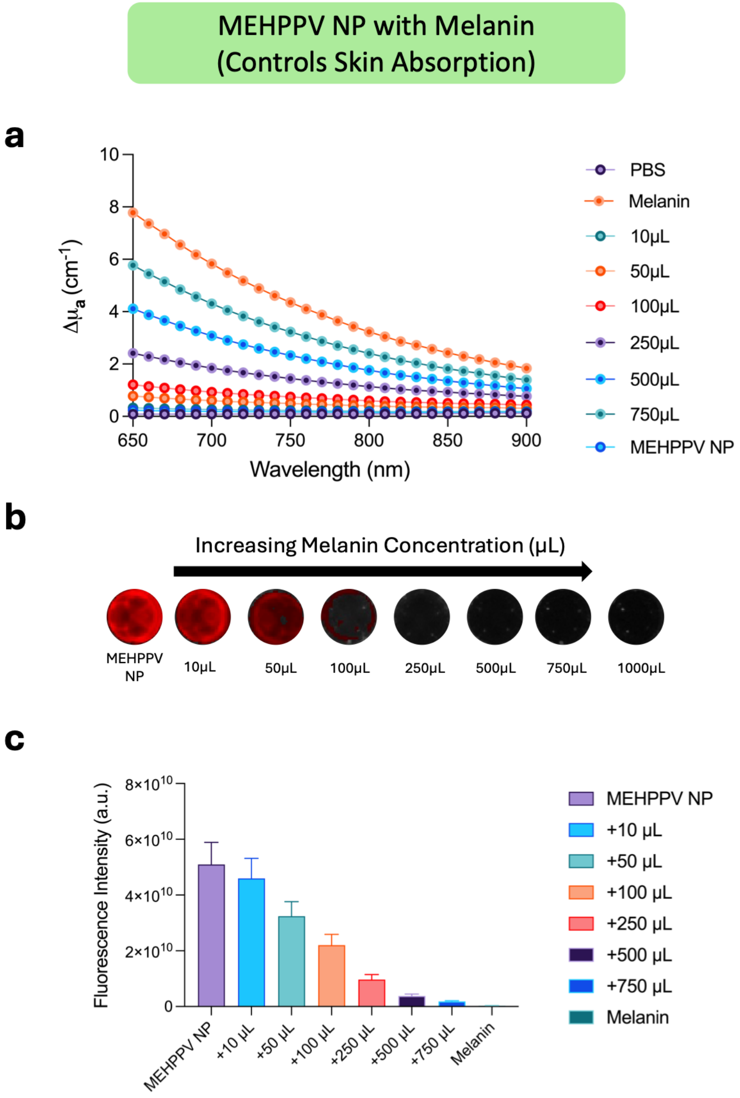


**Figure S12. Effect of the added volume of melanin concentration on the fluorescence of MEHPPV NPs.** (a) Changes in the absorption coefficient ($\Delta\mu_a$) of MEHPPV NPs with increasing volumes of melanin. (b) Representative fluorescence images of MEHPPV NPs in the presence of increasing volumes of melanin (10-1000 μL). (c) Quantitative analysis of fluorescence intensity as a function of added melanin volume. Data are presented as mean ± SD.

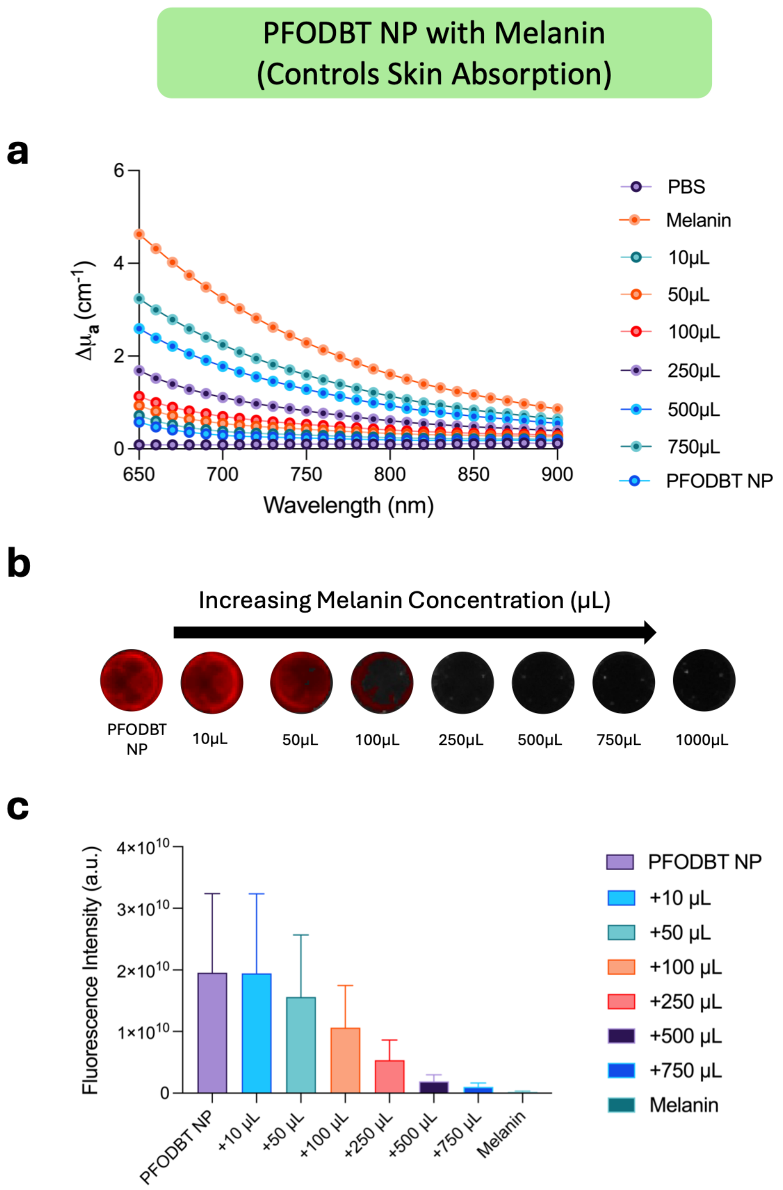


**Figure S13. Effect of the added volume of melanin on the fluorescence of PFODBT NPs.** (a) Changes in the absorption coefficient ($\Delta\mu_a$) of PFODBT NPs with increasing volumes of melanin. (b) Representative fluorescence images of PFODBT NPs in the presence of increasing volumes of melanin (10-1000 µL). (c) Quantitative analysis of fluorescence intensity as a function of added melanin volume. Data are presented as mean ± SD

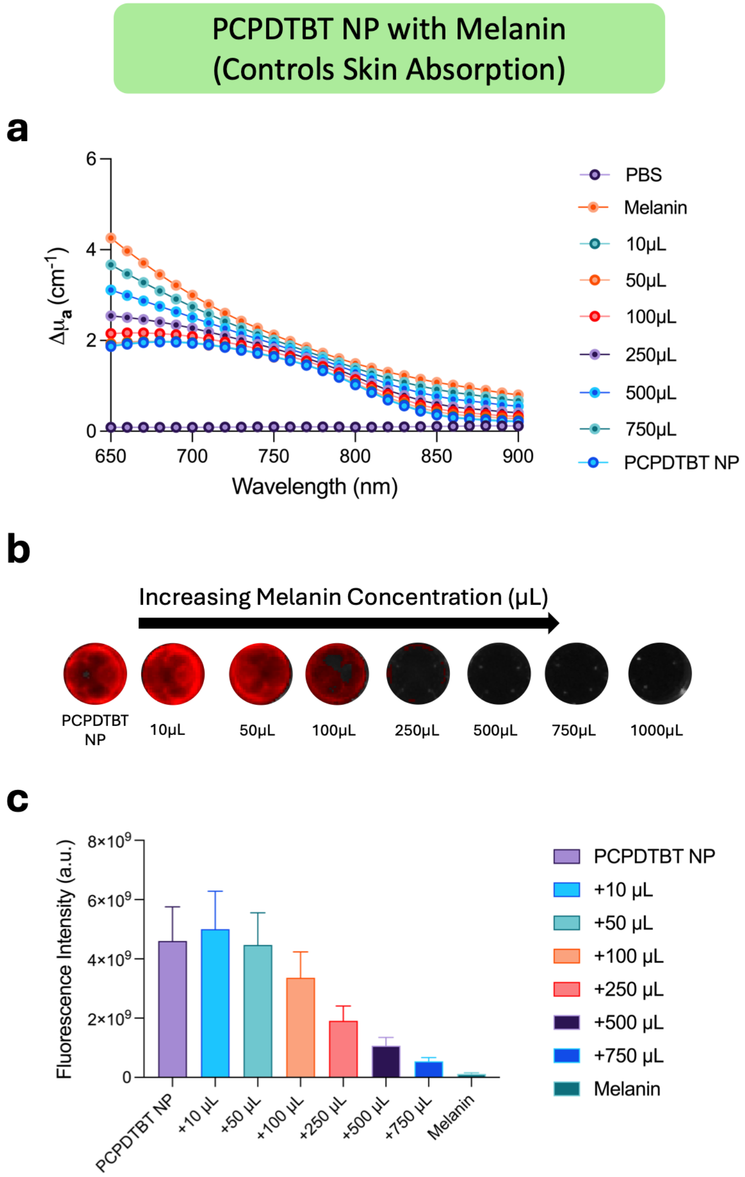


**Figure S14. Effects of the added volume of melanin on the fluorescence of PCPDTBT NPs.** (a) Changes in the absorption coefficient ($\Delta\mu_a$) of PCPDTBT NPs with increasing volumes of melanin. (b) Representative fluorescence images of PCPDTBT NPs in the presence of increasing volumes of melanin (10-1000 μL). (c) Quantitative analysis of fluorescence intensity as a function of added melanin volume. Data are presented as mean ± SD.

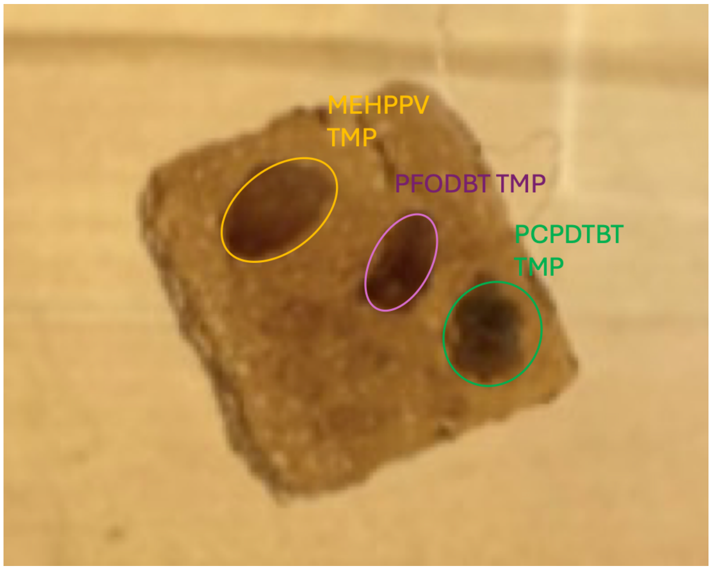


**Figure S15.** Photograph of an intraperitoneal tumor-mimicking phantom containing MEHPPV-, PFODBT-, and PCPDTBT-based TMPs within a skin-mimicking construct.

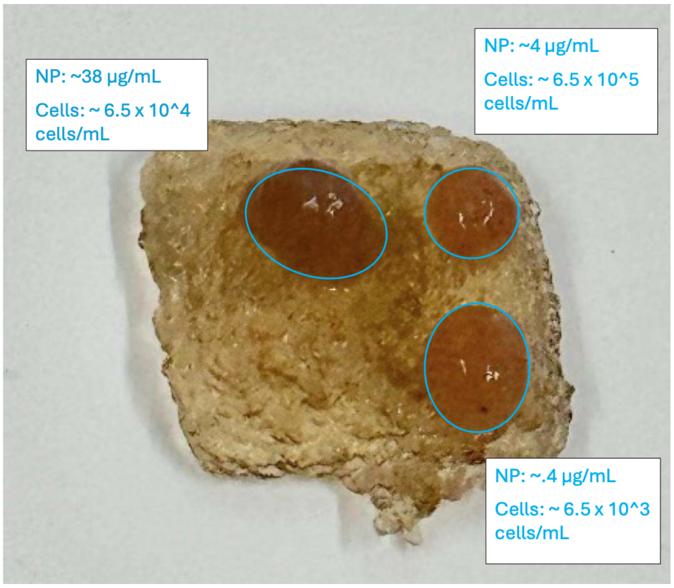


**Figure S16.** Representative photograph of intraperitoneal tumor-mimicking phantoms containing OVCAR-3 tumor cells and PCPDTBT NPs at the indicates concentrations.

**Table S1:** Physiochemical characterizations of NPs.

| Nanoparticle | $D_h$ (nm) | PDI (a.u.) | ZP (mV) |
|---|---|---|---|
| MEHPPV | 194.75±33.16 | 0.0287± 0.03 | -3.726 ± 4.33 |
| PFODBT | 180.3516±83.04 | 0.2120±0.02 | -1.395 ± 7.214 |
| PCPDTBT | 152.522±44.73 | 0.0861±0.10 | -2.900 ± 4.231 |
